# Probing DNA damage sites reveals context-dependent and novel DNA damage response factors

**DOI:** 10.1101/2024.10.23.619792

**Authors:** Richard Cardoso da Silva, Davide C. Recchia, Vincent Portegijs, Maria Cristina Trueba Sanchez, Kelly E. Stecker, Sander van den Heuvel, Tuncay Baubec

## Abstract

DNA damage is a constant threat to genome integrity and function. Diminished capacity for DNA repair is linked to many human diseases, therefore understanding the molecular pathways responding to DNA damage is key for developing novel therapies. Lack of unbiased probes to report DNA damage dynamics and the associated proteins in living cells and animals limit our current efforts to completely understand DNA repair processes. In this study we overcome these limitations by engineering protein probes containing the tandem-BRCT domain of MCPH1, which we show to have a specific affinity for the DNA-damage-associated histone mark γH2AX. We employ these probes to track DNA damage dynamics in living cells exposed to a panel of different genotoxic insults and to visualize programmed double strand breaks during gametogenesis in living animals. We further utilize the binding selectivity of our probe to tether TurboID biotin ligases to chromatin and identify the DNA damage-associated proteome via proximity ligation. By comparing five different DNA damaging agents, we reveal the proteome associated with specific lesions, and identify multiple novel proteins with potential implications in damage response and repair. Among these novel proteins, we characterize the ubiquitin ligase UBE3A, the methyl-binding and proteasome-recruiting protein L3MBTL3, and the spliceosomal factor U2SURP, as previously uncharacterized effectors of DNA damage response. These functional datasets reveal the DNA damage-dependent proteomes and reveal novel insights into DNA damage response.

## Introduction

Genome stability is constantly threatened by DNA lesions triggered by endogenous and exogenous factors (Ciccia and Elledge., 2010). Double Strand Breaks (DSBs), one of the most prevalent and deleterious types of DNA damage, trigger the activation of the DNA Damage Response (DDR). This comprises a set of coordinated signalling pathways that elicit the hierarchical recruitment of repair proteins to DNA damage sites (Dantuma and Attkkum, 2016). In eukaryotic cells, DSB detection and repair occur in the context of chromatin (Kinner et al., 2008). The responses to DSBs substantially rearrange the local chromatin environment through a wide range of chromatin modulators, histone modifying enzymes, histone chaperones, and chromatin remodelers (Price and D’Andrea., 2013; Dabin et al., 2016; Chakraborty et al., 2021). In response to DSBs, the C-terminal tail of histone H2A is phosphorylated. In mammals, the variant H2AX is phosphorylated (γH2AX) by phosphatidylinositol 3-kinase (PI3K)-like protein kinases, including ATM and DNA-PK specialized kinases (Kinner et al., 2008). Phosphorylation of H2AX is not limited to the immediate vicinity of DSBs but spreads to larger chromatin regions influenced by loop extrusion and topologically associated domains (TAD) (Rogakou et al., 1998; Collins et al., 2020; Arnould et al., 2021). γH2AX serves as a signaling platform that guides spatiotemporal recruitment of DNA repair factors to DSBs (Paull TT et al., 2000).

Currently, monitoring of γH2AX distribution, quantification, and kinetics in a spatiotemporal manner is highly constrained by the availability of effective live cell-compatible probes. Apart from some recent advances using nanobodies (Rajan et al., 2015; Moeglin et al., 2021; Trakarnphornsombat and Kimura., 2023), methods employing conventional antibodies directed against γH2AX in fixed/permeabilized cells are more commonly used. Although such approaches have proven very efficient in scoring γH2AX distribution on chromatin, their use as probes to follow the kinetics of DSB formation is very limited. Furthermore, detection of γH2AX in living organisms or other biological specimens that could potentially be employed as a biomarker for drug development, exposure to radiation, as well as for cancer chemo- and radio-therapy trials, has also been hampered by the lack of a specific and efficient probes. Finally, the emergence of whole-genome screens and proteomic methodologies has led to the discovery of numerous proteins involved in DSB repair and signalling pathways (Gupta et al., 2018; Abbasi and Schild-Poulter., 2019; Scott et al., 2021). Yet, the complete repertoire of regulatory proteins triggered by DDR in different cellular or genotoxic contexts is still not known and a comprehensive list is required to understand the underlying repair mechanisms and to identify novel therapeutic targets.

Here, we developed novel DNA damage-specific sensors that allow live monitoring of DSBs in living cells, as well as to identify the protein composition around DNA damage sites. Based on our previous work (Villaseñor et al., 2020), we developed engineered chromatin readers (eCRs) containing selected naturally occurring tandem BRCT domains, and screened for eCRs highly selective for yH2AX. BRCT domains have been previously characaterized as “readers” of phosphorylated yH2AX histones (Mesquita et al., 2010). We show that the stable expression of a specific Microcephalin 1 (MCPH1) tandem-BRCT containing eCR allows scoring of the dynamics of yH2A(X) distribution on chromatin in living cells exposed to different DNA damage insults and in living animals. By fusing this eCR to a promiscuous biotin ligase we could unbiasedly detect the protein-chromatin interaction networks associated with DNA damage sites in cells exposed to different genotoxic agents. This allowed us to identify previously unreported DDR factors, such as UBE3A, U2SURP and L3MBT3, and confirm their role in regulating 53BP1 recruitment to DNA damage sites.

## Results

### Generation and selection of engineered DNA damage sensors

To develop an engineered chromatin reader (eCRs) for detection of DSBs in living cells, we explored the specificity and affinity of DDR proteins that are recruited to yH2AX sites in response to the presence of DSBs. BRCA1 C-terminal (BRCT) domains recognize and bind linear motifs phosphorylated by kinases that are activated following DNA damage (Manke et al., 2003). These evolutionary conserved domains are frequently present in proteins involved in the DDR and are found either as single units or in tandem copies (Mesquita et al., 2010; Woods et al., 2012), separated by a variable linker region (Woods et al., 2012). A database search for mouse BRCT domains resulted in 49 singleton BRCT units grouping in six distinct branches based on amino acid sequence similarity (Figure 1A). We narrowed down our selection to tandem BRCTs from proteins whose localization to sites of DNA damage are well documented. Our final selection comprised tandem BRCT domains present in the following proteins (Figure 1B): MDC1 (BRCT 1-2) (Lou et al., 2006), BRCA1 (BRCT 1-2) (Jeon et al., 2012), PAXI1 (BRCT 5-6) (Jowsey et al., 2004), TP53BP1 (BRCT 1-2) (Ward et al., 2003), MCPH1 (BRCT 2-3) (Wood et al., 2007), DNLI4 (BRCT 1-2) (Grawunder et al., 1998) and TOPBP1 (BRCT 5-6) (Wang et al., 2011). Several of these domains have been characterized as modules responsible for recruiting DDR factors to yH2AX sites (Manke et al., 2003). Amongst them are the tandem BRCTs of the mediator of DNA damage checkpoint (MDC1) that regulates the recognition and repair of DSBs (Lee et al., 2005; Stucki et al., 2005), and of BRCA1 itself, which play major roles in DSB repair (Clapperton et al., 2004; Li and Yu., 2013). The cDNAs encoding the selected tandem BRCTs were stably expressed as GFP-fusion eCRs from a defined site in the mouse ES cell (mESC) genome, as previously described (Villaseñor et al., 2020) (Figure 1B). The selected eCRs displayed stable and homogenous expression, as assessed by measuring eGFP levels by flow cytometry (Figure S1A). Furthermore, eCR expression did not influence cell viability upon induction of DSBs (Figure S1B).

**Figure 1.**
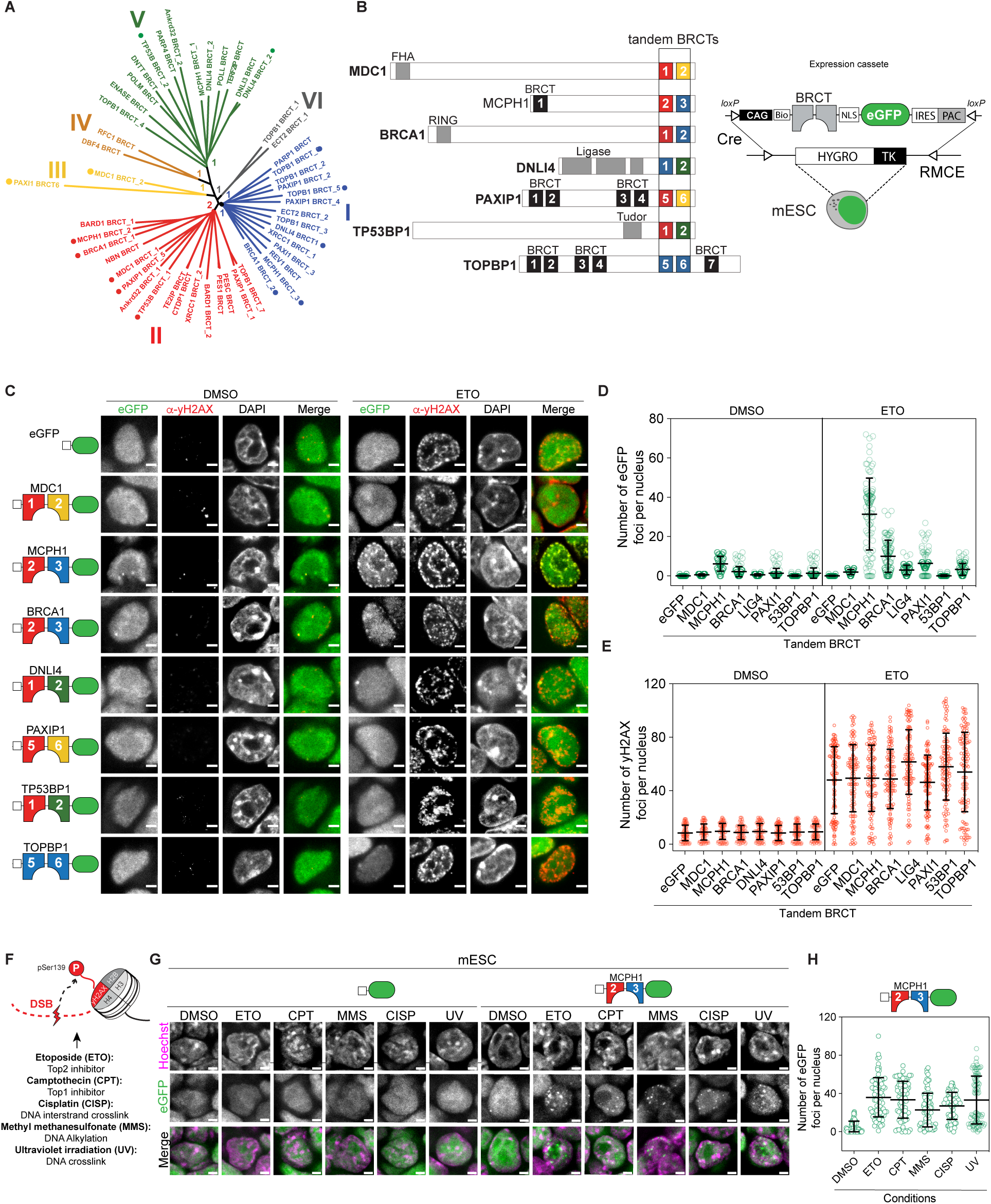
- Tandem MCPH1-BRCT eCR localizes to DNA damage foci marked by yH2AX. **(A)** Unrooted tree illustrating the conservation and relationship between individual BRCT domains. BRCT domains from *Mus musculus* were clustered based on multiple amino acid sequence alignments using the maximum-likelihood method. Support values for main branches (bootstrap test with 1000 replications) are shown. Colors are based on the clustering of singleton BRCTs (I to VI). Numbers shown after the underscore in protein names refer to the relative position of the singleton BRCT in each protein. Dots designate the selected individual BRCT domains used in the rest of this study. **(B)** *Left*, schematic representing the selected mouse proteins containing the tandem BRCT domains domains used in this study. Individual BRCT domains are highlighted and colored based on their clustering from the unrooted tree. *Right:* Schematics of the eCR expression cassette used in mESCs to generate stably expressed BRCTs fused with eGFP. The construct contains a biotin acceptor site (bio), and nuclear localization signal (NLS) under the control of the CAGGS promoter (CAG). RMCE followed by double selection was applied to generate stable integration of the expression construct at a defined site in the mouse genome with the aid of a Cre recombinase. **(C)** Representative immunofluorescence images of fixed mESCs showing the nuclear localization of the indicated tandem-BRCT domains fused to eGFP (green) and their colocalization with yH2AX foci (red). A nuclear eGFP was used as a negative control. Cells were treated with ETO for 1h and stained with antibodies against yH2AX. **(D)** Quantification of the number of eGFP foci per nucleus in mESCs from C (DMSO or ETO-treated). The data represent mean ±SD from at least 100 cells. Comparable results were obtained in two additional independent experiments. **(E)** Quantification of the number of yH2AX foci per nucleus in mESCs from C. Data representation and number of samples according to D. **(F)** Schematic of Histone H2A variant X, phosphorylated at Serine 139 (yH2AX) upon DSB formation triggered by different DNA damage agents. **(G)** Representative images of fixed mESCs expressing MCPH1-BRCT-eGFP or eGFP alone. Cells were treated for 1h with the different DNA damaging agents as indicated, fixed, and imaged right after staining of DNA with Hoechst (magenta). **(H)** Quantification of the number of microscopically discernible eGFP foci per nucleus in mESCs from G. Data representation and number of samples according to D. All scale bars are 5 µM.

To assess the specificity of the selected eCRs for DSB sites, we quantified eCR-GFP foci by immunofluorescence and measured their colocalization with yH2AX foci in mESCs exposed to etoposide (ETO), a drug that induces DSBs through the inhibition of topoisomerase II (Figure. 1C). Amongst the selected eCRs, the tandem BRCT domain of MCPH1 showed the largest number of distinct foci colocalizing with sites of DNA damage marked by yH2AX (Figures 1C-E and S1C). The BRCA1-BRCT-eCR also displayed increased number of foci associated with DNA damage, while the additional eCRs showed reduced numbers of eGFP foci and varying degrees of colocalization to yH2AX sites in presence of ETO (Figures 1C-E). This was not due to different levels of DNA damage in the analysed lines since quantification of nuclear foci revealed comparable numbers of yH2AX signals in treated and control cell lines (Figure 1E). Furthermore, the results obtained in fixed cell preparations were also observed by live cell imaging, excluding potential cross-linking artefacts (Figure S1D).

Although recruitment of BRCT domain-containing full-length proteins to DSB sites has been previously established, our results indicate that the tandem BRCT domain of MCPH1, and to some extent the tandem BRCT domain of BRCA1, functions as a single structural unit and is sufficient for recruitment to yH2AX sites (Figures 1C-D). Surprisingly, the isolated tandem BRCT domains from the well-characterized yH2AX-binder MDC1 (Stucki et al., 2005) did not show a strong association with yH2AX in our assays, suggesting that additional domains present within the full-length MDC1 protein are required for recruitment to DNA damage sites. We confirmed these observations by performing pull-down experiments in mESC lines exposed to Camptothecin and the PARP inhibitor Olaparib (CPT/OLAP), where only the MCPH1-BRCT-eCR but not MDC1-BRCT, co-purified yH2AX (Figures S1E-F).

Given its recruitment to yH2AX sites, we selected the MCPH1-BRCT-eCR and further examined its application as a DNA damage sensor. Besides ETO, damage-induced MCPH1-BRCT foci formation was also observed in mESCs treated with various DNA-damage or replication-stress agents that substantially triggered yH2AX, including CPT, Methylmethanesulfonate (MMS), Ultra Violet radiation (UV), and Cisplatin (CISP) (Figures 1F-H and S1G-H), further confirming that the selected eCR can relocate to sites of DNA damage triggered by distinct agents.

### Functional evaluation of MCPH1-BRCT-eCR interactions with yH2AX

Next, we examined whether individual BRCT domains of MCPH1 are sufficient for the recruitment to DSB sites. We generated mESC lines stably expressing either the BRCT2 or the BRCT3 domain alone and examined their recruitment to yH2AX sites upon treatment of cells with ETO (Figure 2A). Recruitment of both individual BRCTs to yH2AX sites is partially compromised relative to the recruitment of the tandem MCPH1-BRCT (Figures 2A and 2B), further confirming that their association in tandem plays a crucial role for recruitment to DSB sites. To assess the specificity of the selected eCRs, we introduced point mutations known to disrupt the recruitment of full-length MCPH1 to DNA damage foci to the BRCT domains (Wood et al., 2007). We generated cell lines expressing MCPH1 tandem BRCT harboring a single point mutation within the BRCT2 (MCPH1 BRCT2-W706R-BRCT3-WT) or BRCT3 (MCPH1 BRCT2-WT-BRCT3-W815R) domains. Recruitment to ETO-yH2AX sites was strongly reduced in both cell lines, much more so than the recruitment of individual wild type BRCT2 or BRCT3 (Figures 2A and 2B). Furthermore, inhibition of ataxia-telangiectasia-mutated (ATM), one of the kinases that phosphorylates H2AX in the presence of DSBs, significantly reduced formation of MCPH1-eGFP foci in mESCs exposed to CPT/OLAP (Figures S2A and S2B), further confirming the specificity of this recruitment. Altogether, these data strongly suggest that the MCPH1-BRCT eCR is specifically recruited to DNA damage sites by binding yH2AX via its BRCT tandem domain.

**Figure 2.**
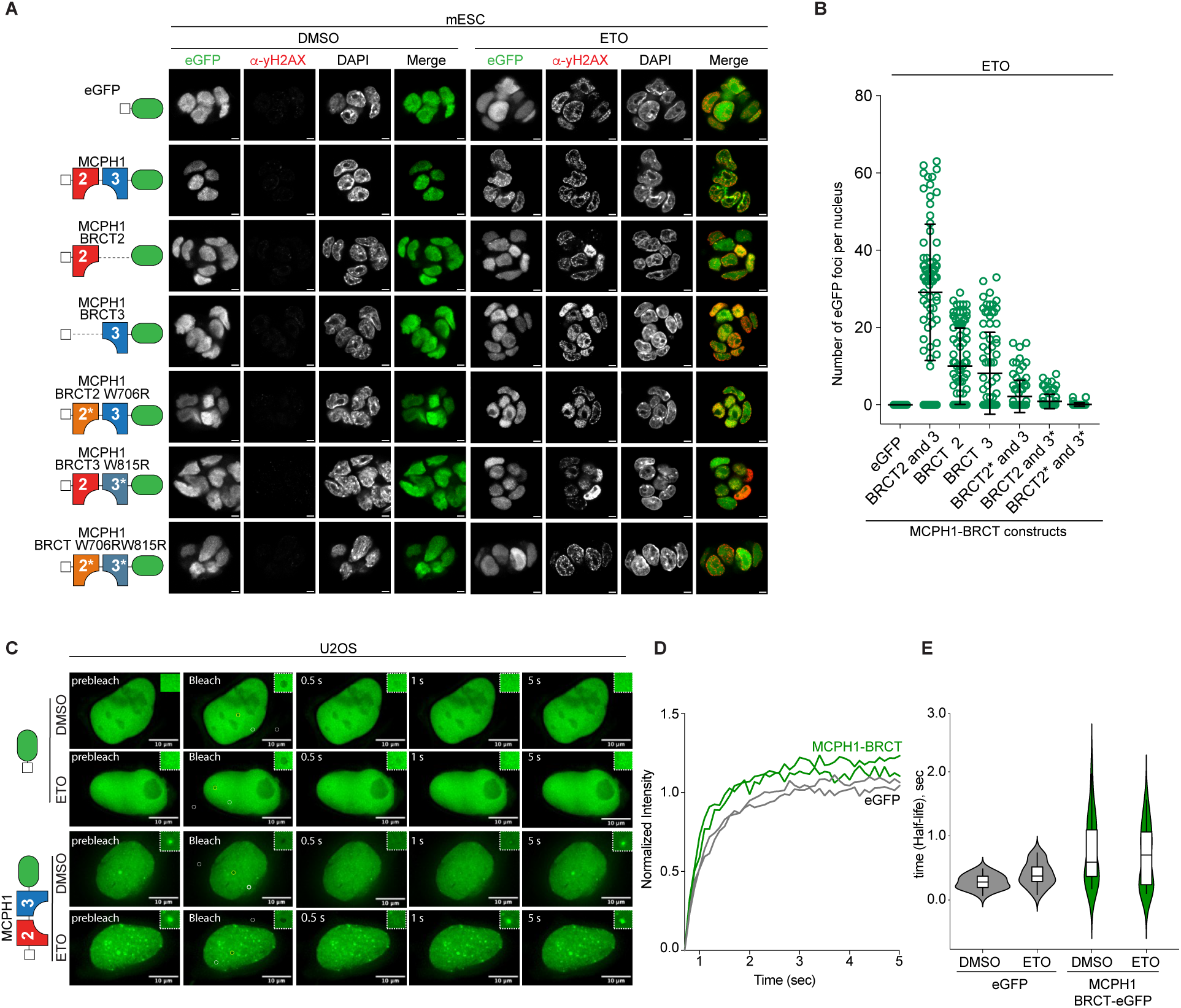
- Specific affinity of the tandem BRCT domain of MCPH1 to yH2AX. **(A)** Representative immunofluorescence images of fixed mESCs stably expressing the indicated constructs (Singleton BRCTs and BRCTs harboring the indicated loss of binding point mutations marked with an asterisk) fused to eGFP. The nuclear localization of eGFP is shown in cells treated as indicated for 1h and stained with antibodies against yH2AX. Scale bars: 5 µM. **(B)** Quantification of the number of eGFP foci per nucleus in mESCs from A treated with ETO. The data represent mean ± SD from at least 80 cells. Two independent experiments showed comparable results. **(C)** Representative images from FRAP experiments performed in U2OS cells stably expressing MCPH1-BRCT fused to eGFP or the nuclear eGFP control. Cells were treated as indicated for 1h. Pre-bleached, bleached, and post-bleached representative images are shown. Zoomed regions of interest (ROI) are depicted on the upper-right side of each image. Scale bars are shown. **(D)** Quantification of FRAP recovery time up to 5 seconds post-bleach. Data shows the mean GFP intensity (pre-bleached and background-normalized ROIS) **(E)** Estimation of the half time of recovery (half life) (*τ*_1/2_), in seconds, from the post-bleach recovery curves depicted in Figure S2L. Data represents mean ± SD from at least 20 ROIS in 20 randomly selected cells per condition. Two independent experiments showed the same results.

mESC harbour a short G1 and G2 phase and a long S phase relative to proliferating somatic cells, and despite displaying a highly active DDR, they also lack a G1 checkpoint (Tichy and Stambrook., 2008). Furthermore, yH2AX sites in mESCs are associated with extensive loading of the single-strand binding proteins RPA32 and RAD51, which majorly confines these cells to repair via Homologous Recombination (Ahuja et al., 2016). Therefore, we aimed to evaluate the specificity and affinity of the eCR MCPH1-BRCT towards yH2AX in a different DDR context. We generated human osteosarcoma (U2OS) cells carrying stably integrated MCPH1-BRCT eCRs (Figure 1B). Two independently derived clones with homogenous eCR expression (Figures S2C and S2D) showed viability kinetics comparable to wild type U2OS cells when challenged with different concentrations of ETO (Figures S2E and S2F). Consistent with the findings obtained in mESC, the MCPH1-BRCT eCR colocalized to yH2AX-marked damage sites after exposure of U2OS cells with ETO (Figure S2G). Moreover, additional microscopy experiments enabled the visualization of discrete nuclear MCPH1-BRCT-eGFP foci upon DSBs induced by a diverse range of DNA damage-inducing drugs (Figures S2H-K).

To gain insights on the dynamics of the MCPH1-BRCT eCR binding to chromatin, we performed FRAP (Fluorescence Recovery after Photobleaching) in U2OS cells and calculated the recovery half-life of MCPH1-eGFP foci from independent nuclei upon exposure to ETO for 1 hour and cells treated with DMSO. The half-life of MCPH1-BRCT-eGFP in ETO damaged nuclei (t-hl = 0.82 sec ±0.71) was nearly identical to undamaged control cells (t-hl = 0.78 sec ± 0.34), suggesting that the binding and diffusion behaviour of MCPH1-BRCT is not affected when chromatin is exposed to DNA damage (Figures 2C-E and S2L). Furthermore, the averaged retention half-life at foci in MCPH1-BRCT cells was only twice as long compared to the half-life measured in cells expressing diffuse NLS-eGFP, which served as non-interacting protein control (t-hl = 0.42 sec ±0.27; t-hl = 0.47 sec ±0.31, undamaged and damaged chromatin, respectively) (Figures 2C-E and S2L). This suggests that MCPH1-BRCT repeatedly binds to and dissociates from chromatin in a dynamic manner.

### MCPH1-BRCT sensors track yH2AX kinetics in living cells and animals

Currently, conventional antibodies directed against γH2AX in fixed/permeabilized cells are very efficient in scoring γH2AX distribution on chromatin. However, their use as probes to follow the kinetics of DSB formation is very limited. We therefore investigated whether the MCPH1-BRCT eCR can be employed as a DNA damage probe to track DSBs in living cells. Live-cell imaging followed by Hoechst staining confirmed the formation of MCPH1-BRCT eCR foci upon induction of DSB by CPT in both mESC (Figures 3A and 3B) and U2OS cell lines (Figures 3C and 3D). Time-lapse confocal microscopy following addition of CPT was conducted in both cell types including eGFP controls over a period of 90 min, capturing images every 5 min (Figures S3A and S3C). Estimation of the number of foci per nucleus revealed a gradual increase in the number of MCPH1-BRCT-eGFP foci already after 15 min following addition of CPT in both cell lines (Figures S3B and S3D). Extended acquisition over a period of approximately 15 hours in mESCs revealed a gradual increase in the number of foci post exposure to CPT, plateauing around 6h, and remaining stable up until ∼ 15h (Figures 3E-F; Videos S1-S4). This was different from results obtained in U2OS cells, where formation of eCR foci peaked at 9h post CPT treatment and decreased over time (Figures 3G and 3H; Videos S5 and S6). This difference is likely due to the absence of a G1 checkpoint which may allow mESC to enter S phase without fully repairing the DSBs (Tichy and Stambrook., 2008), while U2OS cells promote DSB repair before entering S phase. The specificity of the eCR sensor was confirmed by point mutations in both MCPH1-BRCT domains, which significantly reduced the number of fluorescent eGFP foci upon treatment of mESCs and U2OS with CPT. Importantly, the point mutations also reduced the number of MCPH1-BRCT foci in undamaged mESC and U2OS cells (DMSO-treated) attesting the sensitivity of the eCR as a yH2AX readout (Figures S3E and S3F).

**Figure 3.**
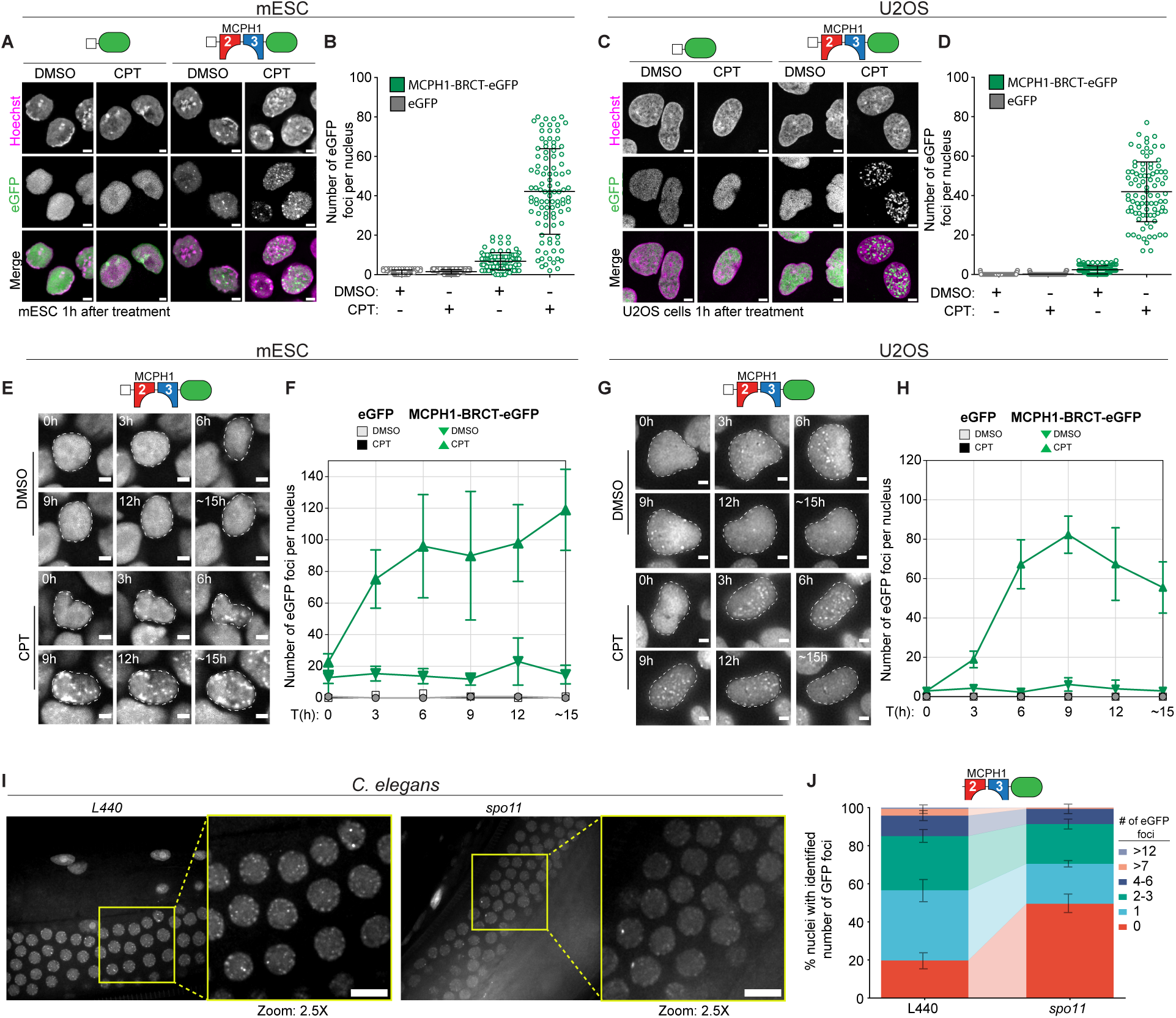
- Using the MCPH1-BRCT eCR to track yH2AX kinetics in living cells and animals. **(A)** Representative live-cell imaging of mESCs stably expressing MCPH1-BRCT-eGFP or eGFP alone. Cells were treated as indicated for 1h and stained with Hoechst (magenta) prior to imaging. Quantification of the number of eGFP foci per nucleus in mESCs from A. The data represent mean ± SD from at least 100 segmented nuclei. Three independent results showed similar results. **(C)** Representative live-cell imaging of U2OS cells stably expressing MCPH1-BRCT-eGFP or eGFP alone. Cells were treated as in B. **(D)** Quantification of the number of eGFP foci per nucleus in U2OS cells from C, measured from at least 100 segmented nuclei. Two independent experiments showed comparable results. **(E)** Representative filmstrips of time-lapse fluorescence microscopy obtained from proliferating mESCs expressing MCPH1-BRCT-eGFP at different time points and imaged after the indicated treatments (from videos S1 and S2). The ROI around selected nuclear eGFP is shown. **(F)** Quantification of nuclear eGP foci from mESCs expressing MCPH1-BRCT-eGFP or nuclear eGFP at different time points. Cells were imaged every 5 minutes right after the addition of the indicated agents and quantification was performed with at least 50 nuclei from two independent time-lapse microscopy experiments using images captured every 3h. **(G)** Representative filmstrips of time-lapse fluorescence microscopy obtained from proliferating U2OS cells expressing MCPH1-BRCT-eGFP and imaged according to E (from videos S5 and S6). All scale bars from A-H are 5 µM **(H)** Quantification of nuclear eGP foci from U2OS cells expressing MCPH1-BRCT-eGFP or nuclear eGFP at different time points. eGFP foci per nuclei were quantitated according to F from at least 50 cells obtained from two independent time-lapse microscopy experiments. **(I)** Representative confocal images of MCPH1::NLS::eGFP in control and *spo-11* depleted live *C. elegans*. 2.5X zoomed images are also depicted. Scale bars: 50 µM. **(J)** Percentage of nuclei with the indicated amount of MCPH1::eGFP foci in wild type and *spo-11* animals. Data represent mean ±SD from 600 to 650 randomly selected nuclei from at least 11 animals.

To further explore the applicability of the MCPH1-BRCT eCR as a damage sensor beyond induced DNA damage in cell culture conditions, we introduced a slightly modified construct expressing the eGFP tagged MCPH1-BRCT eCR into the *C. elegans* genome to monitor programmed meiotic DNA DSBs induced by the topoisomerase-like protein Spo-11 (SPO-11). We detected the presence of distinct MCPH1-BRCT-eGFP foci in the nematode germline which was significantly reduced upon depletion of *Spo-11* (Figures 3I and 3J). Collectively these results indicate that the MCPH1-BRCT eCR is able to capture DSB repair kinetics and can be employed as a damage sensor in living cells and animals.

### Expression of MCPH1-BRCT eCR is compatible with endogenous DSBs in living cells

Having shown that the MCPH1-BRCT eCR is a suitable DNA damage sensor, we wanted to exclude the possibility that it interferes with global DNA repair kinetics and cellular function. Cells stably expressing the eCR showed similar sensitivity to different doses of UV radiation and different concentrations of CPT and ETO, as observed in cells expressing eGFP only, or wild type U2OS cells (Figures S4A-C). Clonogenic survival assays of U2OS cells exposed to UV radiation, further confirmed these results, and suggest that expression of the eCR does not interfere with the ability of cells to grow and form colonies (Figures S4D and S4E).

By monitoring the kinetics of yH2AX in U2OS cells continuously exposed to ETO for 24 hours or pulsed for three hours and then grown in medium without ETO for 24 hours, we could test whether the presence of eCRs affect the ability of cells to recover from DNA damage. Measurement of yH2AX levels by flow cytometry and Western Blot at different time points did not indicate differences between wild type cells or cells expressing MCPH1-BRCT-eGFP or eGFP alone (Figure S4F). In all cases, a partial clearance of yH2AX was observed already after 12h post release from ETO, whilst an almost complete clearance was observed 24h post-release, indicative of completed DNA repair. Levels of yH2AX within the cells continuously exposed to ETO kept increasing and plateaued at 24h (Figures S4G-H and S5A-B). In addition, cell cycle profiling using bivariate DNA/PCNA flow cytometry analysis did not reveal differences in the cell cycle progression of U2OS cells expressing either MCPH1-BRCT-eGFP, eGFP, or wild type cells (Figures S5C-D). Altogether, these results support that expression of the MCPH1-BRCT eCR does not interfere with endogenous repair processes.

### Defining the chromatin-associated protein composition at DNA damage sites

DDR involves a multitude of interactions between proteins and DNA damage sites. Profiling the spatiotemporal changes in the DNA/chromatin-associated proteome surrounding DSBs is essential for our understanding of the DDR. After assessing the affinity of the MCPH1-BRCT eCR towards yH2AX and its feasibility for scoring DSBs in living cells, we aimed to exploit its specific association to damaged chromatin to unbiasedly identify protein-chromatin interaction networks associated with DNA damage sites with the aid of proximity biotinylation. Towards this, we fused the MCPH1-BRCT eCR with the promiscuous biotin ligase TurboID and added a flexible linker to increase the radius of biotinylation labeling (Kim et al, 2016) (Figure 4A). We established U2OS cell lines stably expressing this construct or expressing a nuclear TurboID protein as background control (Figure 4A). Presence of these constructs in U2OS cells leads to increased biotinylation of proteins upon addition of biotin to the culture medium (Figure S6A). Furthermore, we demonstrate that the tagged MCPH1-BRCT eCR efficiently tethers the TurboID ligase to sites of DNA damage marked by yH2AX in ETO-treated U2OS cells, resulting in increased biotin accumulation (Figure 4B). In contrast, nuclear TurboID control does not present any specific binding to chromatin (Figure S6B). We enriched nuclear biotinylated proteins using streptavidin-based affinity enrichment, followed by stringent washing steps, and identified biotinylated proteins using quantitative label-free liquid chromatography followed by tandem mass spectrometry (LC-MS/MS) (Figure 4C). Global analysis of proteins revealed a high correlation among three technical and two biological replicates (Figures S6C and S6D).

**Figure 4.**
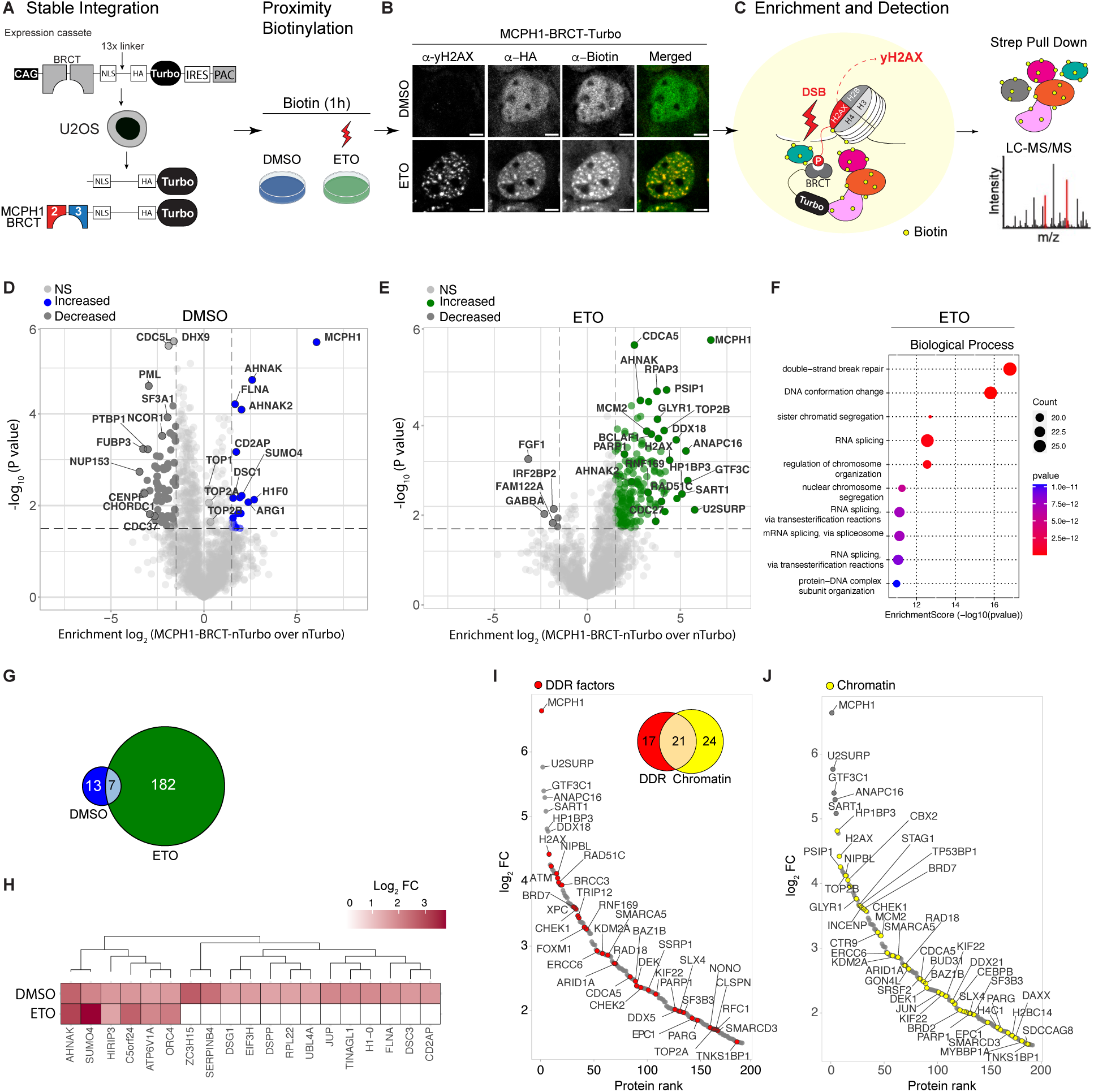
- Probing DNA damage sites with the MCPH1-BRCT eCR reveals the proteome composition associated with DNA damage sites. **(A)** Schematic representation of the ChromID-based proximity biotinylation approach: Left: expression cassette used to generate U2OS cells stably expressing MCPH1-BRCT fused to the TurboID biotin ligase and an HA tag. CAG: CAGGS promoter; NLS: nuclear localization signal. A 13X flexible linker is present between the eCR and TurboID. Nuclear TurboID is used as a negative control. Right: Conditions used in the proximity biotinylation experiments depicted in D-J **(B)** Representative immunofluorescence images of U2OS cells showing the recruitment of MCPH1-BRCT-Turbo to yH2AX sites. Following the co-incubation of cells with biotin and with the indicated agents, cells were co-stained with anti-yH2AX, anti-HA, and anti-biotin antibodies. Scale bars: 5 µM. **(C)** Schematic representations: Left: MCPH1-BRCT-Turbo-dependent biotinylation of proteins in close proximity to DSBs marked by yH2AX. Right: Enrichment of biotinylated proteins by streptavidin pull-downs followed by LC/MS/MS: liquid chromatography coupled to tandem mass spectrometry. **(D and E)** Volcano plot depicting ChromID results. The x-axis represents the log_2_ fold change (FC) and the y-axis represents the -log10 FDR corrected *p*-value determined by a two-tailed *t*-test between MCPH1-BRCT-TurboID over the TurboID-NLS (nTurbo) control performed at the indicated conditions. Selected enriched proteins are highlighted. Light gray denotes proteins not statistically significant. Peptides used to identify MCPH1 matched the tandem BRCT used to generate the DNA damage eCR. **(F)** Bubble chart showing the top ten enriched Biological Process GO terms (based on *P-values*) associated with statistically enriched proteins (MCPH1-BRCT-Turbo/nTurbo, treated with ETO, Log_2_FC>1.5, n=189). The total number of variables from E (n=1769) was used as background. **(G)** Venn diagram showing the overlap of the statistically significant protein hits of MCPH1-BRCT-Turbo after DMSO or ETO treatments. **(H)** Heatmap representation of significantly enriched proteins in MCPH1-BRCT-Turbo datasets from DMSO-treated cells. Shown are LFQ intensities (log_2_-FC). The Log_2_FC values of MCPH1 itself was excluded from the heatmap. Proteins not identified in ETO conditions are shown as white boxes in the heatmap. **(I-J)** Ranking of proteins significantly enriched after ETO treatment based on their log_2_-FC. Classification according to their associated DDR GO term: GO:0006974 (marked in red) and/or chromatin GO term: GO0000785 (marked in yellow). A Venn diagram depicting the overlap between proteins classified by the mentioned GO terms is shown.

Under unperturbed conditions the MCPH1-BRCT proximal proteome encompassed 19 high confidence hits (log_2_ fold change >1.5). Apart from the identification of AHNAK, a protein associated to the p53 Binding protein 53BP1 (Ghodke et al., 2021), most of the proteins identified are general chromatin proteins not directly connected to the DDR, such as Histone H1, the histone chaperone HIRIP3, and ORC4 (Figures 4D and 4H). Upon induction of DSBs with ETO for 1h we identified 189 high confidence DNA damage-associated proteins (Figure 4E). Consistent with the affinity of the MCPH1-BRCT towards DSBs sites, GO terms, such as “double strand break repair”, “DNA conformation change,” and “sister chromatid segregation” were significantly overrepresented in the enriched dataset (Figures 4F, S7A and S7B). Amongst the identified hits, six of them, besides MCPH1 itself, were also identified in unperturbed cells, including AHNAK, HIRIP3 and ORC4, suggesting that these proteins might be basally associated to endogenous yH2AX-DSB sites (Figure 4G-H and S7C; Table S2). We also identified proteins reported in previous studies, including one of the kinases of H2AX, ATM, and classical proteins involved in DNA repair such as 53BP1, RAD51C and BRCA1 (Gupta et al., 2018). Additionally, we also found proteins involved in chromatin remodeling (CHD1, CHD7, CHD8 and SMARCA5) as well as proteins involved in RNA processing that have been previously found in association with the DSB-associated protein Ku, such as DDX5 and SNRNP200 (Abbasi and Schild-Poulter., 2019), (Figures S7C; Table S2).

To further investigate the nature of these hits we clustered the proteins identified in ETO-treated U2OS cells based on their GO classification into two main groups: associated to chromatin (GO:0000785) or DDR (GO:0006974). 38 of the identified proteins classified as DDR, with 21 of them also being associated to chromatin. 24 additional proteins were associated to chromatin only (Figures 4I and 4J). The remaining “non-DDR and non-chromatin” proteins belong to additional classes, including RNA processing, spliceosome, or cell cycle (Figure S7D). Given that not all proteins involved in the DDR are known and that the gene ontology reference curation is likely incomplete, the actual fraction of DDR-related proteins associated to DSBs in our dataset is likely underestimated. Within the DDR-associated proteins we observed a clear enrichment of factors linked to DSB repair, including hits specifically associated with HR and NHEJ, while only a limited number of proteins associated to non-DSB repair such as NER and MMR were identified (Figures 5G and S8H).

**Figure 5.**
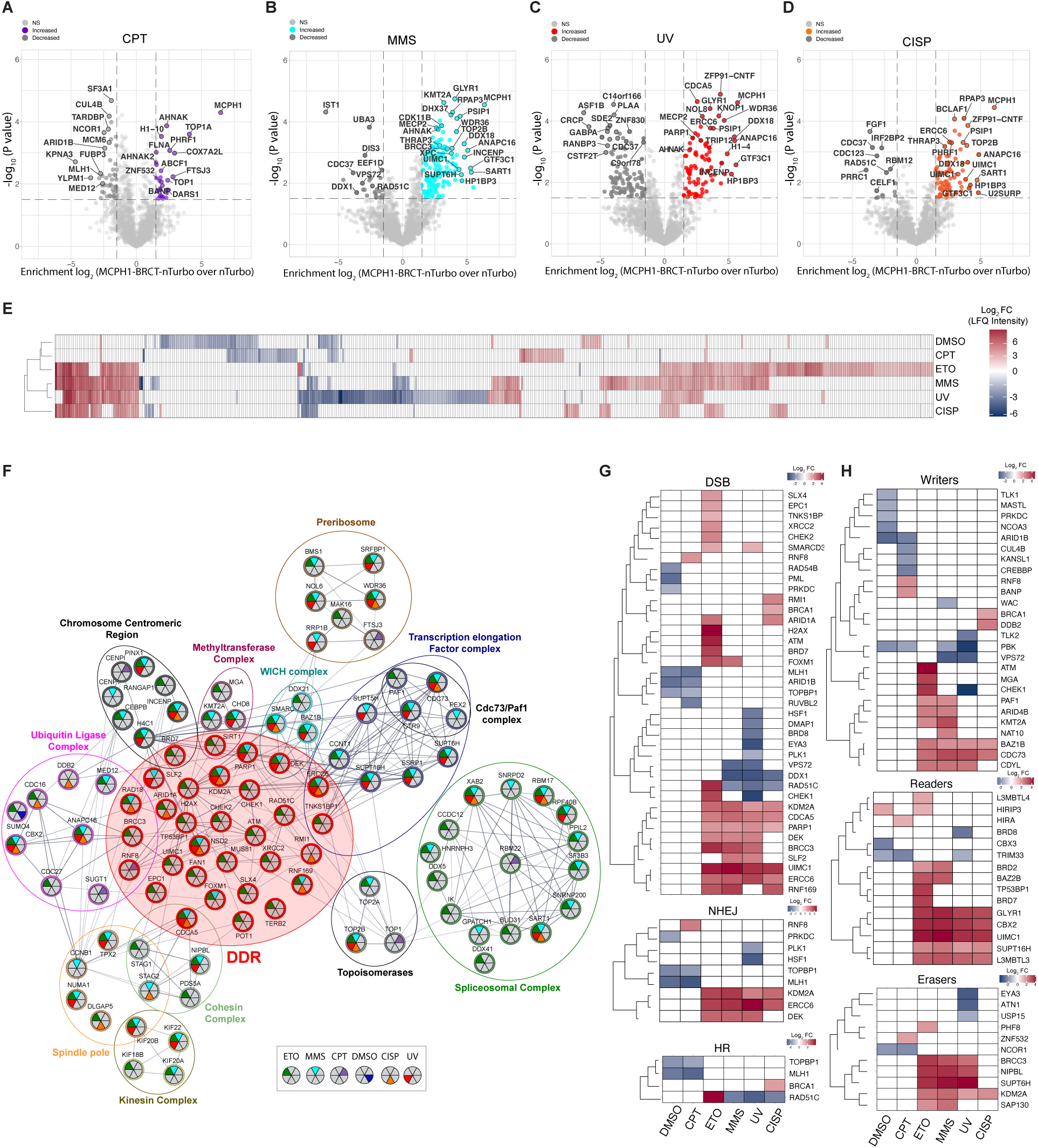
- The proteome composition associated with DNA damage sites is influenced by the genotoxic context. **(A-D)**:Volcano plots showing proteins associated with MCPH1-BRCT-Turbo over nTurbo after treatments with **(A)**: CPT, **(B):** MMS, **(C):** UV and **(D):** CISP. Statistically significant proteins are highlighted. Light-gray dots denote proteins not statistically significant. Selected enriched proteins are indicated. Peptides used to identify MCPH1 matched the tandem BRCT used to generate the DNA damage eCR. **(E)** Heatmap depicting hierarchical clustering of computed LFQ intensities of proteins significantly identified in all the conditions (Log2FC ≤ and ≥ 1.5 of MCPH1-BRCT-Turbo over nTurbo). Proteins not identified in specific conditions are shown as white in the heatmap. The Log_2_FC values of MCPH1 itself was excluded from the heatmap. **(F)** Proteins identified in at least one of the 6 conditions were used to build a network and clustered by the top twelve cellular component GO terms. Proteins associated with the DDR GO term (GO:0006974) were also included in the analyses (red shaded circle). The circled areas show members of protein complexes identified in the network. The names of protein complexes are indicated and individual proteins are shown as nodes. The background color of the protein in the node follows the color attributed to a particular complex. Edges designate physical interactions (experiments) retrieved from the STRING database. Proteins found in different treatment conditions are identified by the color inside the node (see key). **(G and H)** Heatmap visualization of significantly identified proteins (Log2FC ≤ and ≥ 1.5) classified according to different functional categories in the DDR (**G)** and chromatin regulation **(H)**. Proteins were clustered according to their LFQ (log_2_ FC), obtained under different conditions. Proteins not identified in some conditions are shown as white in the heatmap.

### The DNA damage type defines the yH2AX-associated proteome

We next set out to assess differences in protein composition driven by different DNA damage insults. Toward this, we exposed cells to additional DNA damaging conditions (CPT, MMS, UV, and CISP) to obtain a comprehensive view of the context-dependent DDR proteome (Figure S8A). This allowed us to expand the number of identified hits associated with DSB sites to a total of 284 unique proteins significantly enriched across all the different conditions and datasets (Figures 5A-D). Exposure of U2OS cells to CPT, MMS, UV, and CISP, which trigger DSBs via distinct molecular mechanisms relative to ETO, revealed a diverse range of proteins either uniquely present or shared among all the conditions (Figures 5A-E and 4D-E). Exposure of cells to ETO allowed us to identify a higher proportion of hits relative to other conditions (Figures 5E and S8B), which can also be explained by the increased levels of MCPH1-BRCT foci elicited by ETO in relation to other conditions (Figures S2J and S2K). While only a few significant hits were shared amongst all the datasets, most of the common hits were observed in cells exposed to MMS, where unique hits were shared with either ETO or UV (Figures S8B and S8C). Consistent with their effect in triggering DSBs via DNA replication stress, GO terms such as “regulation of chromosome organization” and “chromosome segregation” were significantly overrepresented in cells exposed to MMS and UV (Figures S8E and S8F).

We noticed that most proteins identified upon exposure to ETO, UV or MMS were not identified in cells exposed to CPT or CISP (Figures 5A, 5D and 5E). These displayed specific subsets of proteins (Figure 5E, S8B and S8C) and distinct GO terms such as “associated to microtubule-cytoskeleton organization”, “involved in mitosis” as well as “spindle orientation” following CPT, and “sister chromatid and chromosome segregation” following CISP exposure (Figures S8D and S8G). The specific DNA damage signaling pathways inflicted by these agents are not fully understood and despite inducing DSBs by distinct mechanisms, a synergism between both agents (CPT and CISP) through the increase of DNA interstrand cross-links have been previously described (Kigawa et al, 1999; Fukuda et al, 1996). Thus, it is possible that DSBs triggered by these treatments are spatially distinct in comparison to the other tested conditions. Importantly, the unique identification of TOP1 but not TOP2 upon exposure to CPT (Figure 5A) is in agreement with its role in inhibiting TOP1 by forming stable topoisomerase-I DNA-cleavable complexes. This was also true for ETO-treated cells which were enriched for TOP2A and TOP2B but not for TOP1 (Figure 4D).

To further visualize the cellular distribution and functional relationship of the detected proteins in light of the different DNA damage insults, we created an interaction network of all proteins based on interaction scores obtained from the STRING database, and grouped them by their top “cellular component” GO terms (Figure 5F). In this network we further highlighted the DNA damage-dependent enrichment of individual proteins. The clustering revealed several proteins belonging to distinct regulatory pathways and complexes. Importantly, the network revealed that these complexes interconnect to a larger DDR-centric hub containing many proteins known to be involved in the DNA damage repair and response (Figure 5F). Next, we clustered all identified DDR components across the different treatments to quantify their distribution in a damage-context-dependent manner. We observe that canonical proteins involved in DSB repair such as ERCC6, UIMC1 and PARP1 are present in all conditions, except for CPT and DMSO (Figures 5G and S8H). Other proteins, such as RNF8, RAD51 and BRCA1 were uniquely enriched upon either CPT, ETO or CISP treatments, respectively. This highlights the applicability of our DNA damage sensor to characterize repair pathways triggered by different genotoxic agents, which display unique and shared DDR proteins.

### Chromatin regulators frequently associate with DNA damage sites

Besides known components of the DDR machinery, the network analysis revealed a broad range of proteins and functional complexes known to be associated with chromatin and transcriptional regulation, as exemplified by the recruitment of proteins associated with heterochromatin, such as MECP2, CBX2 and UHRF1 as well as factors associated with regions of active transcription, as illustrated by the enrichment of CDC73 and PAF1 (Figure 5F and Table S2). This is consistent with the interconnection of the DNA damage machinery with chromatin regulation and gene transcription (Gaillard and Aguilera, 2016). Moreover, several centromere and kinetochore-associated proteins were also enriched, which is in line with the localization of several DDR factors to these chromosomal structures (Pageau and Lawrence., 2006; Di Paolo et al., 2014; Saayman et al., 2003). To quantify the damage-context distribution of chromatin regulators, we further clustered the hits into different chromatin regulator classes and measured their enrichment across the different treatment conditions. Chromatin readers such as BRD7 and BRD2 were preferentially located at DSB sites upon ETO treatment, whereas CBX2 and GLYR1 were identified in all the conditions, except for DMSO and CPT (Figure 5H). Several Zinc finger proteins have been implicated in DSB repair and we have identified some of them, for example, ZNF281, herein specifically identified upon exposure to CPT, has been shown to be recruited to DSB sites to facilitate NJHEJ (Nicolai et al., 2019). ZC3H18, identified in all the conditions but DMSO and CPT, is implicated in the activation of BRCA1 (Kanakkanthara et al., 2019). Our analysis also identified numerous other zinc finger proteins not previously reported to be associated with the DNA damage response, ZFP91, ZNF532, ZNF608 and ZC3H15 (Figure S8I).

Cells respond to DSBs via increased accumulation of cohesin near the damaged site (Litwin et al, 2018). Our profiling of different DNA damage conditions enabled the identification of some components of the cohesin complex, including CDCA5, NIPBL, STAG1 and STAG2 (Figure 5F). CDCA5, which stabilizes cohesin on DNA (Ladurner et al., 2016) was significantly identified in all conditions, except for DMSO and CPT, while the cohesin loading protein NIPBL was identified in ETO, MMS and UV conditions. Cohesin containing the more abundant STAG2 subunit has been shown to be essential for chromatid cohesion at centromeres while cohesin containing the less abundant STAG1 subunit seems to be essential for chromatid cohesion specifically at telomeres (Canudas and Smith., 2009; Remeseiro et al., 2012). While the STAG2 variant subunit of cohesin was enriched in MMS and CISP, the STAG1 variant was solely enriched in ETO conditions (Figure 5F and S9A). To test if the differential recruitment of STAG1 and STAG2 to DNA damage types is indeed dependent on the damaging agent, we measured their association with yH2AX following treatment with either ETO or MMS. Immunofluorescence microscopy in unperturbed U2OS cells revealed the expected localization of both STAG1 and STAG2 within the nucleus. Exposure of cells to either ETO or MMS slightly changed the localization of STAG1 within the nucleus revealing a minor overlap of yH2AX foci with STAG1 in the ETO treatment in relation to MMS. Staining of STAG2 revealed an increase in punctate foci in both ETO and MMS conditions and an increased association of STAG2 to DSB sites marked by yH2AX following exposure to MMS (Figures S9B and S9C). Global levels of STAG1 and STAG2 in perturbed cells were comparable to untreated cells (Figure S9D).

### Identification of U2SURP, UBE3A and L3MBTL3 as novel regulators of DDR

To further demonstrate the ability of our eCR to identify novel proteins surrounding DSB sites, we selected three hits that were significantly enriched upon DNA damage but have previously unreported roles in DDR: U2SURP, a subunit of the U2 small nuclear ribonuceloprotein (snRNP), L3MBTL3, a histone methyl-lysine binding protein and UBE3A, a ubiquitin-protein ligase E3A. While UBE3A was only significantly identified in datasets of U2OS cells exposed to ETO, L3MBTL3 and U2SURP were also identified in cells exposed to MMS and CISP (Table S2). Knock-down of these proteins by small interfering RNA (siRNA) in U2OS cells (Figure S10A) allowed us to measure the effect of these proteins on DDR. Towards this, we scored yH2AX and 53BP1 foci by microscopy and assessed the cell viability and cell cycle status of cells recovering from damage insults following gene knock-down of these selected candidates. Depletion of either UBE3A or L3MBTL3 significantly affected the recruitment of 53BP1 to DSBs marked by yH2AX sites following CPT and ETO treatment relative to control cells, resulting in a reduction of visible 53BP1 foci. This decrease was even more pronounced in UBE3A-depleted cells (Figures 6A, 6B, and S10B). Levels of spontaneous or induced DSBs marked by yH2AX were not affected and increased to the same levels under both conditions relative to DMSO (Figures 6C and S10C). In contrast to UBE3A and L3MBTL3, knockdown of U2SURP led to the accumulation of discernible 53BP1 foci in the nucleus already in unperturbed DMSO conditions (Figures 6A and 6B). However, this accumulation was only associated with a minor increase in the global levels of yH2AX and number of yH2AX foci per nucleus (Figures 6A, 6C, and S10C), suggesting a role for U2SURP in preventing DNA damage-independent accumulation of 53BP1. Following CPT and ETO exposure, U2SURP-depleted cells showed a decrease in the number of yH2AX foci relative to control cells, while the number of 53BP1 foci remained elevated (Figures 6A, 6B, 6C, S10B and S10C). Importantly, knockdown of these proteins did not change the global levels of 53BP1 (Figure S10D), suggesting a previously unreported role for these proteins in regulating 53BP1 localization.

**Figure 6.**
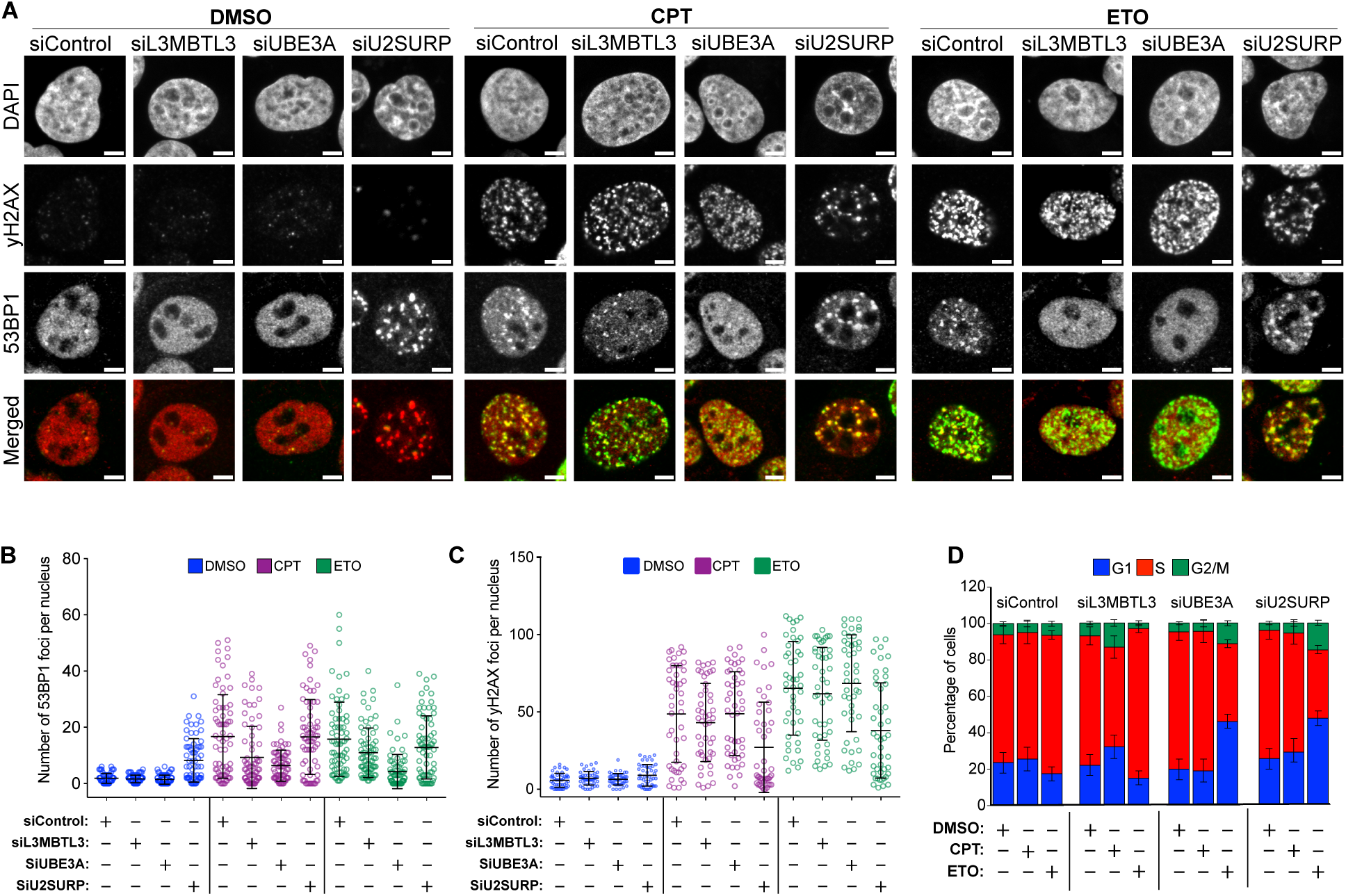
- Depletion of newly identified factors associated with DNA damage sites reveals their effect in the DDR. **(A)** Representative immunofluorescence images of wild-type U2OS cells transfected with the indicated siRNAs for 48h. Following siRNA-mediated depletion of the indicated proteins, cells were treated as indicated for 1h and co-stained with anti-yH2AX (green) and 53BP1 (red) antibodies. All scale bars: 5 µM. Cells from A were used for: **(B)** Quantification of the number of 53BP1 foci per nucleus at the indicated conditions following siRNA treatment for 48h. **(C)** Estimation of the number of yH2AX foci per nucleus at the indicated conditions following siRNA transfection for 48h. Data represent mean ±SD from at least 40 cells. For B and C, comparable results were obtained in two additional independent experiments. **(D)** U2OS cells transfected with the indicated siRNA for 48h were exposed to the indicated treatments for 1h and washed with fresh medium. After 12h their cell cycle distribution was analyzed by flow cytometry. The relative distribution of cells in G1, S, and G2/M is shown. Data represents mean ±SE of three independent experiments.

To test the consequences of the observed changes in 53BP1 recruitment and yH2AX levels upon knockdown of these factors, we measured the viability and cell cycle progression of cells in the presence of CPT and ETO. Depletion of L3MBTL3 showed an effect on cell cycle progression in CPT conditions, whilst the viability of cells in both CPT and ETO conditions was comparable to that of control cells (Figures 6D and S10E-G). Interestingly, knockdown of UBE3A affected the cell cycle progression of cells after exposure to ETO but not to CPT or DMSO (Figures 6D and S10G). This is in accordance with UBE3A being significantly enriched in datasets of cells treated with ETO (Table S2). However, exposure to both CPT and ETO resulted in reduced viability in absence of UBE3A (Figures S10E and S10F). Similar to UBE3A, depletion of U2SURP affected cell cycle progression following ETO treatment, and markedly reduced the viability of cells in treated and untreated conditions (Figures 6D, S10E and S10F).

## Discussion

While many efforts to detect yH2AX in living cells have been made (Rajan et al., 2015; Moeglin et al., 2021; Trakarnphornsombat and Kimura., 2023), current methods to monitor its distribution rely mainly on end-point measurements in cells subjected to fixation/permeabilization and antibodies. Although such methods allow the quantification of yH2AX foci over time, tracking the early kinetics of DSBs in a spatiotemporal manner is limited by the lack of an effective compatible probe. Tagging of endogenous full-length proteins such as 53BP1 (Kilic et al., 2019), or expression of a truncated version, has been used as a probe for DSBs in living cells (Dimitrova et al., 2008). However, recruitment of repair proteins to DSBs can be influenced by the specific repair pathway choice, and therefore preclude their use as a neutral DSB reporter in living cells. In this study we present a novel DNA damage sensor to track DNA damage and provide a comprehensive identification of proteins associated with DNA damage sites in living cells. Toward this, we established engineered chromatin readers (eCRs) selective to yH2AX by screening selected tandem BRCT domains from well-characterized proteins associated with DNA damage response. These eCRs were introduced to mouse stem cells, as previously described (Villaseñor et al., 2020), and assessed for their recruitment to DNA damage sites marked by yH2AX. Amongst the selected candidates, the tandem BRCT domain of MCPH1 (BRCT2 and BRCT3 linked by a small unstructured linker) exhibited the highest affinity towards DSBs. Although single BRCT domains have also been previously shown to interact with yH2AX phosphopeptides (Sun et al., 2017), our data indicates that single BRCT versions of the tandem MCPH1 BRCT were not sufficient to interact with yH2AX sites efficiently - in line with the model where recognition of yH2AX is achieved when the phosphorylated residue is sandwiched by two separate (Shao et al., 2011).

By using several quantitative and functional approaches we have validated the feasibility of the tandem BRCT domain of MCPH1 to cytologically monitor spatiotemporal dynamics and kinetics of DNA DSBs. Microscopy and Flow cytometry analysis indicated that the presence of MCPH1-BRCT on chromatin was detected shortly after the induction of DSBs in the form of discrete nuclear foci and was resolved upon removal of the drug that triggered DSBs without detectably interfering with global DNA repair. Importantly, the evolutionary conservation of H2A phosphorylation at DNA damage sites (Osakabe and Molaro., 2023) allowed us to apply our DNA damage sensor to detect meiosis-specific DSBs in the *C. elegans* germline. These findings highlight the applicability of our eCR to follow the spatiotemporal resolution of DSBs and gain significant insights into DSB kinetics in different model systems ranging from cell lines to living animals. Furthermore, we show that the eCR performs well in both fixed and living cells and can be used as an antibody alternative in standard chromatin immunoprecipitation and immunofluorescence microscopy protocols.

Finally, to identify the proteins associated with DNA damage sites, we fused the MCPH1-BRCT domains to the promiscuous biotin ligase TurboID, resulting in the biotinylation of proteins localised in the proximity of yH2AX. Our analyses confidently captured specific DDR proteins, highly consistent with previously published literature (Gupta et al., 2019; Kratz et al., 2023; Abbasi and Schild-Poulter., 2019) and many novel factors previously not associated with DDR. Importantly, the sensitivity of our assay allowed us to discriminate proteins associated with DNA damage triggered by different genotoxic agents, providing a comprehensive characterization of damage-specific response pathways. Notably, our analysis also identified many non-DDR proteins, associated with chromatin, transcription, RNA processing and splicing. These associations are in agreement with repair of DSBs taking place in the context of chromatin and previously identified chromatin-associated proteins linked to DNA repair (Sigismondo et al., 2023), as well as studies showing the accumulation of specific splicing factors to sites of damage (Beli et al., 2012; Xin-Juan Fan., 2013), which may play roles in regulating the repair process as well as in the coordination of splicing decisions (Tresini et al., 2015; Shkreta and Chabot., 2015; Tanikawa et al., 2016; Fan et al., 2021).

Among the splicing factors identified here, U2SURP, a subunit of the U2 small nuclear ribonucleoprotein complex, is a novel candidate that plays a role in DDR. We show that depletion of U2SURP leads to DNA damage-independent accumulation of 53BP1 foci and a reduction in yH2AX foci in presence of DNA damage. How this splicing factor influences DDR remains to be identified, but potential pathways could involve direct or indirect regulation of 53BP1 stability. Intriguingly, U2SURP, together with CHERP and RBM17, which have also been identified in our datasets (Table S2), have been previously shown to functionally interact and reciprocally regulate protein stability (De Maio et al., 2018). In addition, splicing deficiency leads to deregulation of the E3 ubiquitin ligase RNF8 (Pederiva et al., 2016), which is responsible for ubiquitination of damaged chromatin to promote accumulation of DNA repair factors (Poulsen et al., 2012). The E3 ubiquitin ligase RNF8 was only detected in cells exposed to CPT (Figure 5F and 5G), while other E3 ubiquitin ligases such as RNF169, that negatively regulates 53BP1 (An et al, 2018), CDC27, and E3 ligases whose role in DSB response are currently unknown, such as UBE3A, were also uniquely enriched in cells exposed to DNA damaging agents (Table S2). In the case of UBE3A, we show that absence of this E3 ligase affected recruitment of 53BP1 to DNA damage sites without decreasing its overall abundance, indicating a potential role in deposition of ubiquitin on chromatin surrounding DNA lesions. Finally, we find the methyl-lysine reader L3MBTL3 to similarly influence 53BP1 localization to DNA damage sites. L3MBTL3 binds H4K20me2 (Lindsey I James et al., 2013) and could play a role in modulating 53BP1 binding to this mark, as previously demonstrated for another histone methyl-lysine binding protein, L3MBTL1 (Klara Acs et al., 2011). In addition, recent work has implicated L3MBTL3 in recruiting the CRL4-DCAF5 E3 ubiquitin ligase complex to methylated target proteins based on their amino-acid sequence similarity to methylated histone peptides (Leng F et al., 2018; Zhang C et al., 2019).

While this set of novel proteins with potential involvement in DNA damage repair highlights the power of our approach to identify novel regulatory proteins, the probes developed here will be of great relevance beyond the presented study. We expect the sensors to be applicable to a diverse set of experiments ranging from the identification of proteins associated with DNA damage in wide range of living organisms exposed to genotoxic damage, reliable reporters of DNA damage in biomedical and clinical settings, as well as recombinant proteins that replace antibodies, enabling future studies in the DNA damage field.

## Materials and methods

### Selection of BRCT domains

All singleton BRCT domains from *Mus musculus* (Figure 1A) were manually retrieved from Uniprot. Tandem BRCT domains, comprising two or more closely located BRCT domains linked by a variable linker region were manually retrieved and their association in tandem were further confirmed based on the literature (Mesquita et al., 2010; Leung et al., 2011). Aminoacid sequences of singleton BRCTs were aligned using MAFFT (Katoh et al., 2002) (with the FFT-NS-I iteractive method and default parameters). Amino acid sequences of the selected tandem BRCTs were also aligned using MAFFT using the E-INS-I iteractive method, recommended for sequences with conserved domains and long gaps. Maximum-likelihood unrooted phylogenetic trees were inferred from multiple amino acid sequence alignments with PhyML version 3.0 with the WAG model (Guindon et al., 2010) and edited with the Interactive tree of Life tool (iTOLv5) (Letunic and Bork., 2021). Supporting values for the inferred tree were generated using BOOSTER using 1000 Bootstraps replicates (Lemoine et al., 2018).

### Molecular cloning

The selected mouse tandem BRCT domains (Figure 1B; Table S1) were synthesized by Integrated DNA Technologies (IDT) based on available Uniprot annotations and assembled according to (Villaseñor et al., 2020) into the Recombinase-mediated cassette exchange (RMCE) targeting vector partbit-v6 by Gibson assembly. The final construct harbored the reader domain N-terminally tagged with a biotin acceptor in frame to a cassette containing an NLS, an eGFP followed by the puromycin-N-acetyltransferase gene expressed through an Internal Ribosome Entry site (IRES). Single MCPH1 BRCTs, BRCT2 and BRCT3 were cloned into the same constructions without the linker. Point mutations in the BRCT2 and 3 were based on (Wood et al., 2020). For the TurboID constructs, reader domains were assembled into the RMCE targeting vector parbit-v9 that harbour the same structure as parbit-v6 except that the assembled reader is N terminally tagged with a 1X FLAG and is linked to an HA-TurboID via a 13X linker (Butz et al., 2022). Expression of all constructs is driven by the constitutive CAG promoter.

### CRISPR/Cas9-mediated genome editing

The he423 [peft-3::MCPH1 BRCT::NLS::eGFP(GLO)] allele in the strain SV2569: he423 [peft-3::MCPH1 BRCT::NLS::eGFP(GLO)] was generated using the homology-directed repair of CRISPR/ Cas9-induced DNA double-strand breaks, for which the reagents were micro-injected into the gonad of young adults. The allele was generated using the Alt-R CRISPR/Cas9 system (IDT), as previously described (Ghanta et al., 2021). As a repair template, MCPH1 BRCT was amplified from a gBlock (IDT) using primers with 5’ SP9 modifications (IDT). MCPH1 BRCT was inserted into a pre-existing allele by generating double-stranded breaks using two sgRNAs. To verify the edit, the insertion was PCR amplified and sequenced by Sanger sequencing.

### RNA-mediated interference (RNAi)

For RNAi of *C. elegans*, bacterial cultures of *E. coli* HT115 containing L4440 empty vector or vector with genomic gene inserts were grown overnight, induced with 1 mM IPTG for 1 hour, 5 times concentrated and seeded onto NGM plates containing 12.5 μg/ ml Tetracycline, 100 μg/ml Ampicillin and 2 mM IPTG. RNAi feeding initiated from L4 stage and the number of germline foci were analyzed in late l4/young adult animals of the following generation. RNAi clones for *spo-11* were obtained from the Ahringer (Fraser et al., 2000) database and sequence verified.

### Mammalian cell Culture and cell lines

Mouse embryonic stem cells (HA36CB1) were cultured as previously described (Villaseñor et al., 2020) and stable mESC lines were obtained by RMCE (Baubec et al., 2013). Briefly, constructs were co-transfected using Lipofectamine 3000 reagents (Thermo Fisher Scientific), with a Cre-recombinase expression plasmid into RMCE-competent mESC (HA36CB1), followed by selection with 3mM ganciclovir (Selleckchem) and 2 μg/ml puromycin (InvivoGen). Individual clones were picked and homogenous and stable expression of proteins was confirmed by flow cytometry or Western blotting. Human U2OS cells (kindly provided by Dr. Susanne MA Lens, University Medical Center Utrecht, Netherlands) were cultured in Dulbecco’s modified Eagle’s medium (DMEM), supplemented with 1% penicillin–streptomycin antibiotics and 10% FCS (Thermo Fisher Scientific). For generation of stable U2OS expressing either eGFP or HA-TurboID proteins, parental wild type U2OS cells were seeded in six-well plates at 0.1 x 10^6^ cell/ml and lipo transfected using Lipofectamine 3000 reagent (Thermo Fisher Scientific) for 48h with PvuI-linearized plasmids (Parbitv-6 or Parbitv-9 construcrs). Following incubation for 48h, cells were selected with complete media containing 2 μg/ml puromycin (InvivoGen, ant-pr-1) for 10 days and maintained with 0.5 μg/mL Puromycin. Individual clones, obtained through limited dilution of resistant polyclonal cell population in 96 well plates, were expanded and constructs integration were confirmed by FACS or Western Blotting. Where indicated, the following compounds were used at the indicated final concentrations, unless stated otherwise: ATMi KU-55933 (10 μM, Selleckchem, S1092), Camptothecin-CPT (25nM, Enzo Life Sciences - ALX-350-015), Cisplatin-CISP (10μg/mL, Millipore - 232120), Etoposide-ETO (20μM, MedChemExpress - HY-13629), Methyl methanesulfonate-MMS (0.25mM, Sigma-Aldrich - 129925), Olaparib-OLAP (1μM, Sigma-Aldrich - SML3705). For UV irradiation, culture media was removed and cells were washed once with DPBS. Cells were then subjected to UV-C (254 nm) at different mj/cm2 rates using the GS Gene Linker UV Chamber (Bio-Rad). Cells were then immediately incubated with fresh and warm media.

### siRNA transfections

Individual siRNA transfections were performed using Ambion Silencer Select siRNAs using Lipofectamine RNAiMAX (ThermoFisher Scientific), according to the manufacturer’s instructions. The following Human-Silencer Select siRNAs (Thermofisher) were used at a concentration of 20 nM: siL3MBTL3 (s39037), siUBE3A (s14604), siU2USURP (s23621). For ATM depletion, 40nM of siRNA (s1708) was used. The Silencer Select Negative Control number 1 (Thermofisher, 4390843) was used as a non-targeting control.

### Flow cytometry

For measuring eGFP levels in live mESC and U2OS cell lines, cells were resuspended in DPBS containing 2% FCS and 1mM EDTA and immediately analyzed on a FACSCanto II or FACSymphony A1 Cell Analyzer (BD Biosciences). For yH2AX measurements in U2OS cells, 1×10^6^ cells were collected, resuspended in PBS and permeabilized with PBS/0.1% Triton X 100/1% BSA for 10 min on ice. Cells were then fixed in ice-cold methanol for 3 min at -20°C. Cells were spun down and pellets resuspended in PBS. Cell were blocked with 1% BSA in PBS for 1h and then incubated with anti-yH2AX-Alexa Fluor 647 antibody (1:200) (Biolegend, 613408) or Anti PCNA-PE conjugate (1:200) (Invitrogen, 12-910-42). Cells were then washed and resuspended in DAPI staining solution (0.1% (v/v) Triton X-100-1mg/mL DAPI (Sigma-Aldrich, MBD0015). Alternatively, cells not incubated with conjugated antibodies were fixed with 70% ice-cold ethanol at 4C for at least 30 min, washed with DPBS, and directly incubated with a DAPI-stained solution. Samples were acquired on a FACSCanto II or FACSymphony A1 Cell Analyzer (BD Biosciences) and data were analyzed using FlowJo software (version 10.9, Tree Star). The proportion of cells in each cell cycle stage was assessed using FlowJo with the Watson-pragmatic model, which fits the Gaussian DNA distribution curves to the stages of the cell cycle or by gating the proportion of cells using the DNA content coupled to S phase distribution (PCNA staining). Quenching of eGFP fluorescence with Methanol fixation was controlled using single staining samples and fluorescence compensation to avoid potential bleedthrough of fluorescent signals.

### Western blotting

To measure the levels of eCRs, 10 μg of crude nuclear extracts, prepared as previously described (Manzo et al., 2017), were resolved by SDS-PAGE, transferred to nitrocellulose membranes (Amersham, GE10600001) and blocked in 5% wt/vol non-fat milk in TBS-0.1% Tween-20. Membranes were then probed with the following primary antibodies overnight at 4°C: anti-eGFP (Abcam, Ab290) anti-HA (Abcam, ab9110), anti-TP53BP1(Abcam, ab172580), anti-SA1 (STAG1), (Bethyl - A302-579A), anti-SA2, (STAG2) (Bethyl - A300-159A) and anti-Lamin-B1 (Santa Cruz Biotechnology, sc-6216). Detection was done with species-specific horseradish peroxidase-conjugated secondary antibodies and Pierce. Peroxidase IHC Detection Kit (Thermo Fisher Scientific). For detection of biotin levels, 20 μg of crude nuclear extracts were resolved in SDS-PAGE gels as above, blocked with 5% wt/vol BSA in TBS-0.1% Tween-20 and probed with Streptavidin-Alexa790 (or with anti-Biotin, (Invitrogen, #31852). To measure yH2AX levels, acidic Histone Extracts were obtained as described in (Villaseñor et al., 2020). Briefly, cells were lysed with NETN buffer (20 mM Tris pH 8, 0.5% (vol/vol) NP-40, 100 mM NaCl and 1 mM EDTA pH 8), supplemented with 1X protease inhibitor cocktail (PIC) (Roche) and dithiothreitol (DTT) (Sigma-Aldrich). Cell lysates were centrifuged and histones were extracted by resuspending pellets with 0.2N HCl overnight. The acidic histone extract was centrifuged and histone extracts were then neutralized with 1N NaOH. 3μg of extracts was resolved by SDS-PAGE. Western Blot and detection was performed as above with the following primary antibodies: anti-yH2AX Phospho Ser. 139 (Biolegend, 613402) and anti-H3 (Abcam, ab1791). Nitrocellulose membranes were stained with Ponceau-S (Sigma).

### GFP Pull-downs

mESCs (wild type, eGFP control, MCPH1-BRCT and MDC1-BRCT fused to eGFP), treated with CPT/OLAP for 1h were harvested and resuspended in nuclear isolation buffer (NIB) (15 mM Tris pH 7.5, 15 mM NaCl, 60 mM KCl, 5 mM MgCl_2_, 1 mM CaCl_2_, 250 mM sucrose, 0.03% NP40) and incubated for 30 min at 4°C on a roller. Cells were harvested at 3200g for 10 min at 4°C and supernatant was discarded. Nuclei were resuspend in 2 pellet volumes of RIPA buffer (25 mM Tris pH 7.5, 150 mM NaCl, 1% NP40, 1% Sodium deoxycholate (NaDOC), 0.1% SDS) supplemented with 1X PIC (Roche) and 1 mM DTT (Sigma-Aldrich). Nuclei were incubated at 4°C on a roller for 30 min, sonicated for 5 cycles (30 sec ON/30 sec OFF) in the Bioruptor (Diagenode) and centrifuged at 14000 x g for 2 min at 4°C. Protein concentration was determined with the detergent-compatible Bradford assay kit (Pierce, 1863028). For GFP Pull downs, 10 μl of GFP-trap magnetic beads (Chromotek, gtma-20) was washed according to the manufacturere’s protocol and incubated with 250 μg of Nuclear Extract. 0.5 μl of Ethidium Bromide (10 mg/ml) was added to each pull down. Pulldown was performed by rotating the beads with the nuclear extracts for 90 min at 4°C. The beads were bound to a magnetic rack and washed with 1 mL of RIPA Buffer repeated three times. Beads were then bound to the magnetic rack and the supernatant was removed. Beads were resuspended in 4X SDS-sample buffer (Laemmli) and analysed by western blotting with anti-eGFP (Ab290), anti-yH2AX (Biolegend, 613402) and anti-Lamin-B1 (Santa Cruz, sc-374015-B10) as described above.

### Immunofluorescence of fixed cells

mESC and U2OS cells were grown on sterile coverslips (0.13–0.16 mm – Ted Pella) coated with 0.2% gelatine (mESCs) or uncoated (U2OS cells). For (Figure S9A), cells were grown on 12-well removable chambers (Ibidi, 81201). mESCs and U2OS cells were fixed in 3.7% formaldehyde (Sigma-Aldrich, 252549) or 2% methanol-free formaldehyde (Thermo scientific, 28908), respectively in 1X PBS for 10 min at Room Temperature (RT) and washed three times with 1X PBS. Cells were then permeabilized with 0.5% Triton X-100 (Sigma-Aldrich) for 5 min on ice followed by 10 min at RT. Cells were blocked at RT for 1 h with blocking buffer, 5% BSA (Sigma-Aldrich) dissolved in 0.1% PBS-Tween 20. All primary and secondary antibodies were diluted in filtered blocking buffer. Primary antibody incubations were performed overnight at 4°C in a humid chamber. Secondary antibodies were incubated for 1h at RT. Following incubation with different antibodies, the coverslips were washed three times with 1X-PBS-0.1% Tween-20 and once with 1X PBS. Next, coverslips were mounted using antifading mounting medium containing DAPI (Vectashield, H2000). For experiments in (Figures 1F and S2J), cells, grown on coverslips as above, were fixed with 2% methanol-free formaldehyde (Thermo Fisher Scientific, 28908) in PBS for 15 min at RT. Cells were then rinsed in PBS and washed twice with PBS for 5 min at RT. Cells were stained with Hoechst (Thermo Fisher Scientific, 62249) for 10 min at RT and washed three times for 5 min with 1X PBS. Coverslips were mounted on glass slides with 4 μl Mowiol-based mounting media (Mowiol 4.88 in Glycerol/TRIS). The following primary antibodies were used: anti-Biotin (mouse, Sigma-Aldrich - 3737373, 1:500), anti-yH2AX (mouse, Biolegend - 613402, 1:1000), anti-TP53BP1(rabbit, Abcam - ab172580, 1:500), anti-HA (rabbit, Abcam - Ab9110, 1:1000), anti-SA1 (STAG1) (rabbit, Bethyl - A302-579A, 1:2000) and anti-SA2 (STAG2) (goat, Bethyl - A300-159A 1:5000). The following secondary antibodies were used: Alexa-Fluor 488 Goat anti-Rabbit IgG H+L (Thermo-Fisher Scientific, A11034, 1:500), Alexa-Fluor 488 Goat anti-mouse IgG H+L (Thermo-Fisher Scientific, 2465113, 1:500), Alexa-Fluor 568 Goat anti-mouse IgG H+L (Thermo-Fisher Scientific, A11004, 1:500), Alexa-Fluor 647 Goat anti-Rabbit IgG H+L (Thermo-Fisher Scientific, A21244, 1:500), Alexa-Fluor 488 Chickent anti-Goat IgG (Thermo-Fisher Scientific, A21467, 1:500), Alexa-Fluor 568 Donkey anti-Goat IgG (H+L) Cross-adsorbed Secondary Antibody (Thermo-Fisher Scientific, A11057, 1:500). Images of fixed cells were acquired with a Zeiss LSM 700 confocal laser scanning microscopy (Biology Image Center, Utrecht University, Netherlands) with a Plan-Apochromat 63X 1.4 NA oil immersion objective (Carl Zeiss, Germany) using 405, 488, 555 and 647 nm laser lines. Images consist of a z-stack of 4-12 planes at 0.37 µm intervals. In Figures S1E, S2A, 3E, 3G, S3A, S3B, S3E and S3F, Images were obtained using a DeltaVision RT Core System (Cell Microscopy Core, UMC Utrecht, the Netherlands) with a 60X objective (Olympus), equipped with Coolsnap HQ + Photometrics EMCCD. Images were deconvolved using the SoftWoRx software 6.1.l Overlap of the emission spectra (bleedthrough) was corrected by automatic and manual adjustments of multiple channels, by using negative control (unstained cells) and by performing sequential scanning.

### Live-cell image microscopy

1.5 × 10^4^ mESCs or 2.5 × 10^4^ U2OS cells expressing either eGFP tagged eCRs or eGFP control cells ∼16h-20h before imaging. Unless stated otherwise, mESC cells were seeded in 8-well Ibidi chambers (Ibidi, 80826) prior coated with 0.2% gelatin. U2OS cells were seeded in uncoated 8-well Ibidi chambers (Ibidi, 80826) or 35 mm Quad-µ-Dishes (Ibidi, 80416). The next day, cells were treated with CPT or DMSO. For one-time images, live mESCs and U2OS cells were stained with Hoechst-33342 (Thermo Fisher Scientific, 62249) for 10 min in the dark right after treatment, and covered with specific mESC or U2OS phenol-red free DMEM (Thermo Fisher Scientific, 31053028). For time-lapse microscopy, environmental controllers were set up so that environmental chambers were stably operating under 5% CO_2_, at 37°C under 80% relative humidity prior to incubation of ibidi chambers. Ibidi chambers containing the cells and phenol red-free DMEM without CPT (concentration of 10 nM for experiments shown in Figures 3E-H) or vehicle control, were carefully transported to environmental chambers. Immediately after positioning of cells, establishment of data points and additional imaging parameters, half of the media was replaced with media containing the desired drug concentration/vehicle (kept at 37°C) and the ibidi chamber was carefully closed. After re-adjusting the image positioning and ensuring that the image fields remained in focus, time-lapse was immediately started. Images were acquired at 5-min intervals in a series of 2-µm-spaced z-sections on a DeltaVision RT imaging system - GE Healthcare equipped with a heated 37°C chamber, photometrics Cascade II EM-CCD camera and four detection channels (Cell Microscopy Core, UMC Utrecht, Netherlands), Images were captured using an Olympus 60X-142 NA-UPLSApo/O oil objective and exposure time setting to 50 ms. Images were deconvolved using the SoftWoRx software 6.1.l and/or z-projected (maximum intensity) using Fiji (ImageJ 64-bit; Version 2.00-rc-54/1.51 h). Images of mESC (Figures 3E and S3A) were acquired at 5-min intervals in an Olympus ScanR Screening System (ScanR Image Acquisition 3.01) (Center for Microscopy and Image Analysis, UZH, Switzerland) equipped with an inverted motorized Olympus IX83 microscope, a motorized stage, IR-laser hardware autofocus, a fast emission filter wheel, a CellVivo incubation system and a Hamamatsu ORCA-FLASH 4.0 V2, high-sensitivity digital monochrome scientific cooled sCMOS camera. Images were captured using a 40x Phase PLAPON (NA 1.42), oil objective. A single z-plane was selected for quantitative image series analysis and representative display.

### Live cell imaging of *C. elegans*

Imaging of *C. elegans* was performed by mounting the larvae on a 5 % agarose pad in 20 mM tetramisole solution in M9 (0.22 M KH2PO, 0.42 M Na_2_HPO4, 0.85 M NaCl, 0.001 M MgSO4). Spinning disk confocal imaging was performed using a Nikon Ti-U microscope equipped with a Yokogawa CSU-X1 spinning disk using a 60X-1.4 NA objective, and a Prime BSI Express Scientific CMOS camera. Spinning disk z-stacks were acquired using MetaMorph Microscopy Automation & Image Analysis Software.

### Fluorescence recovery after photobleaching (FRAP)

For FRAP assays, U2OS cells were grown on 35 mm Quad-µ-Dishes (Ibidi, 80416) and medium containing either ETO or DMSO control was added as described above (live cell imaging). Images were acquired on a Leica Thunder microscope with TIRF and infinity scanner equipped with a prime 95B sCMOS with CO_2_ and temperature control. A circular region of interest (ROI) with 0.34 μm surrounding the selected foci, the nucleus and the background was established using the LasX (Metamorph) software. ROIs were bleached for 55 sec (using a 100x objective with a numerical aperture (NA) of 1.4, a 488 nm laser, 100% intensity). Post-bleach fluorescence was monitored in a time-lapse for a total of 15 sec with a time interval of 0.100 sec. Fluorescence intensities (mean per pixel) were saved for each ROI and used to determine the diffusion coefficients and t_1/2_ half-time of recovery using MATLAB (frap_abalysis, version 1.0.0.0).

### Image processing and data analysis

Image serial stacks were generated and processed with Fiji (ImageJ 64-bit; Version 2.00-rc-54/1.51 h) and appropriate z-project planes were then selected for further image analyzes. For representative images, projection images were selected and processed using Fiji. CellProfiler (Broad Institute, version 4.2.6) (Stirling et al., 2021) was used to estimate the number of yH2AX, TP53BP1 or eGFP foci in fixed cells of mESCs and U2OS cells. Every image was corrected for illumination biases using the “Rescale Intensity” module prior to image analyses. For mESCs, images were also processed with a Gaussian filter to remove the foreground signal. Nuclei (DAPI or Hoechst stained) were segmented with the Otsu threshold and watershed modules to produce a binary image. Segmented nuclei were further identified using the IdentifyPrimaryobjects module. Subsequently, the resulting images were analyzed by a watershed algorithm with varying thresholds to detect foci outlines, depending on the cell line, using the “IdentifySecondaryobjects” function. Drug treatment did not bias the determination of parameters. Missegmented nuclei and nuclei within large colonies from mESCs were manually assessed and discarded from the analyses. The number of eGFP foci per nucleus from time-lapse microscopy of live eGFP-tagged mESCs and U2OS cells was estimated with the Imaris Software (Oxford Instruments, version 10.0.1). Firstly, a surface was created based on the GFP signal background. Generated surfaces were further segmented through a manual histogram-based thresholding. Missegmented nuclei, mitotic cells nuclei and nuclei whose signal was oversaturated (based on assessment of out-of-range pixels) were further discarded. The number of foci per nucleus (green channel) was estimated using the “Spots” function from the Imaris software. Colocalization coefficients (Mander’s correlation), based on pixel intensities of (Regions of interest generated through DAPI segmentation) yH2AX, STAG1, STAG2 and eGFP signals were obtained using the coloc2 plugin from fiji (ImageJ 64-bit; Version 2.00-rc-54/1.51 h). Whenever required, cells were equally adjusted for brightness and contrast. Quantification of the number of eGFP foci per nucleus in C. elegans was done according to (Toraaason et al., 2022) using Imaris Software (Oxford Instruments, version 10.0.1). Individual nuclei were identified as surface objects based on eGFP signal background through manual thresholding. Poorly segmented nuclei were excluded from the analyses. eGFP foci were then identified using the Spots function. To circumvent the lack of a marker to identify DSB transition zones, nuclei were randomly selected in different regions of multiple animals.

### Cell viability Assays

Cell viability assays were performed using the CellTiter-Glo Luminescent Cell Viability Assay, (Promega, G7570), according to the manufacturer’s instructions. Briefly, 5000 mESCs or 7500 U2OS cells were seeded onto black-walled and flat-bottom 96 well plates (Greiner). For mESCs, plates were coated with 0.2% gelatine prior to seeding. Cells were incubated at different time points with different concentrations of DNA damage agents or UV-radiated and allowed to recover at 37°C and 5% CO_2_. Cells treated with DMSO, or non-UV-radiated cells were used as control. Prior to measuring, the same volume of freshly prepared CellTitter-Glo reagent was added and plates were incubated for 2 min on an orbital shaker followed by 10 min incubation at RT. Plates were measured using the luminescence microplate reader CLARIOstar Plus (BMG Labtech). The raw data generated for control treatments, set to 100%, was used to estimate the percentage of viable cells.

### Clonogenic assays

For colony survival assays, U2OS cells were seeded at single-cell density onto 6-well plates overnight (1500 cells per well). On the next day, cells were either UV irradiated and allowed to recover under normal growth conditions, changing the medium every 2 days, or treated with CPT or ETO for 24h, washed three times with 37°C U2OS medium and allowed to recover. After 10 days, medium was removed and cells were rinsed with PBS. Cells were then fixed with 2mL of fixation solution (Methanol:Acetic acid 3:1) and incubated for 5 min at RT After removing the fixation solution, colonies were stained with 0.5% crystal violet (VWR, 20.846.361) prepared in 20% methanol for 15 min. Plates were washed extensively with water and allowed to dry overnight. The number of individual colonies as well as the signal intensity of the whole plate normalized to the background (region on plates devoid of colonies) was estimated with Fiji and values were used to calculate platting efficiency and surviving fraction relative to non-UV-radiated/DMSO treated cells. Clonogenic survival was estimated based on control plates (set to 100%).

### RNA extraction and RT-qPCR

Total RNA was isolated 48h after siRNA transfection using the RNeasy Plus Mini Kit (Qiagen, 74134) according to the manufacturer’s instructions. The integrity of RNAs was assessed by RNA screen Tapestation analysis (Agilent, 5067-5576). cDNA was generated from 1μg RNA using Superscript III reverse transcriptase with oligodT-20 (Thermofisher, 18080044). Relative expression of selected targets was measured by real-time quantitative PCR (RT-qPCR) using Power SYBR Green PCR master mix (Thermofisher, 4367659) on a CFX-Connect Real-Time PCR detection system (Bio-Rad). Relative fold gene expression was analyzed using the Double Delta Ct method (2^-ΔΔCt). The *C*q (Ct) values of the selected gene were subtracted from the housekeeping gene (*GAPDH*). This value was put in the power of 2 and this was also done for the S.D. Values from siControl were set to 1. The oligo sequences from Humans, spanning exon-exon junctions, are as follows:

*GAPDH* (NM_001256799.3):

FW: 5′-AATGGGCAGCCGTTAGGAAA-3′, RV: 5′-GCCCAATACGACCAAATCAGAG-3′; *L3MBTL3* (NM_001007102.4):

FW: 5′-TCACATGAAGCCAGAGGTGC-3′, RV: 5′-AGGGCAGACATGGAATTGGG-3′; *UBE3A* (NM_001354505.1):

FW: 5′-GGCGTTTTCAAGGTTTTTGGC-3′, RV:5′-GGCTCTTACAGATTTTTAACCAAGC-3′; *U2SURP*: (NM_001080415.2)

FW: 5′-ATCCCAACAGAAAGGAATTTGC-3′;

RV: 5′-TTGGCCCTTCACGTACAACA-3′.

### Preparation of nuclear extract for TurboID-proximity ligation

Extraction of nuclear proteins were generated as previously described in (Butz et al, 2022) with some modifications. Briefly, 6×10^6^ U2OS cells were seeded on 15-cm plates and cultured to 70% confluency. Fresh medium containing the selected DSB agent and 50 μM D-Biotin (Thermo Fisher Scientific, B20656) were added and incubated for 1h. All the subsequent steps were either performed at 4°C or on ice. Cells were scraped off from plates with ice-cold PBS, centrifuged at 100 x g for 7 min and swelled on ice with 2mL of nuclear extract buffer 1 (NEB1; 10 mM HEPES, pH 7.5, 10 mM KCl, 1 mM EDTA, 1.5 mM MgCl_2_,) containing 1 mM DTT and 1X PIC (Roche) for 10 min. Pellets were collected by centrifugation at 2,000 x g for 10 min. The supernatant was then removed and the pellet gently resupended in 1mL of ice-cold NEB1. Cells were homogenized using a pre-chilled loose (type-A) Dounce pistil (10 strokes) in 1mL NEB1. Nuclei were collected by centrifugation at 2,000 × g for 10 min, resuspended in 500 μl of NEB1 containing 12μl of 25U Benzonase (Merck, 71206) and rotated for 3h at 4°C. Nuclei were collected as above, resuspended in 500 μl of NEB2 containing 450 mM NaCl (NEB2; 20 mM HEPES, pH 7.5, 0.2 mM EDTA, 1.5 mM MgCl_2_, 20% glycerol, 1 mM DTT and 1× PIC). Nuclei were broken and homogenized with a tight (type-B) pre-chilled Dounce pistil (10 strokes), vortexed and rotated for 1h. Cell debris was removed by centrifugation at 13.000 rpm for 15 min before adjusting the salt concentration to 150 mM NaCl by dropwise addition of NEB2 without NaCL and adjustment of nuclear extract with NP40 to 0.3%. After checking protein concentration with the Qubit Protein Assay kit (Thermo Fisher Scientific, Q33211), the same volume adjusted with immunoprecipitation buffer (IPB; NEB2, 150 mM NaCl, 0.3% NP40, 1 mM DTT, 1× PIC) and concentration of nuclear extract was used per IP.

### Streptavidin Pull Downs, on bead digestions and peptide desalting

10μl of Streptavidin Sepharose High-Performance beads (Cytiva, 17511301) was used per pull down. Beads were washed twice with 1 mL ice-cold RIPA buffer supplemented with 1 mM DTT) and 1X PIC (Roche); inverted 10X and collected at 2000 x g for 2 min at 4°C. Beads were then resuspended in 92 μl of RIPA buffer per pull down and 95μl of beads/IP were collected in pre-chilled protein low binding eppendors (Thermo Fisher, 3448). After discarding the supernatant, the nuclear extract was added and incubation was carried out overnight at 4°C under rotation. Next day, beads were collected and washed (as above) once with ice-cold RIPA buffer, two times with high-salt buffer (HSB; 50 mM HEPES pH 7.5, 1 mM EDTA, 1% Triton X-100, 0.1% NaDoc, 0.1% SDS, 500 mM NaCl, 1 mM DTT and 1× PIC), once with DOC (250 mM LiCl, 10 mM Tris pH 8.0, 0.5% NP-40, 0.5% deoxycholate, 1 mM EDTA, 1 mM DTT and 1× PIC) and four times with ice-cold TE buffer with 1mM DTT (without supplementing with PIC). After collecting the beads by centrifugation, residual TE buffer was carefully aspirated from the beads with the aid of 30G needles. Next, beads were resuspended in 50 μl of elution buffer (2 M Urea, 100 mM Tris pH 8, 10 mM DTT) followed by incubation on a thermo shaker at 1400 rpm for 20 min at RT. Chloroacetamide (CAA) was then added to a final concentration of 50 mM and samples were incubated in the dark for an additonal 10 min. After adding 250ng of Trypsin (Promega, V5113), samples were incubated for 2h in a shaker. Supernatants were then transferred to another low protein binding eppendord and beads were trypsinized another time and collected as above. Supernatants were then combined and digested with 100 ng of trypsin overnight at RT. The next day, digested peptides were acidified by the addition of 5% (v/v) LC-MS grade Formid Acid (FA) (Merck, 5.33002.0050) and purified on C18 columns (Stagetips) (Pierce, 87784) C18 columns were washed once with LC-MS grade methanol (Thermo Fisher Scientific, 47192), two times with solution A (60% LC-MS grade Acetonitrile (SUPELCO, 1.00029.1000), 0.1% FA) and two times with solution B (0.1% FA) prior to loading the samples. After collecting the peptides in low-binding protein Eppendorf, C18 columns were washed once with buffer B. Columns were then stored at 4°C and washed once prior to elution with solution A. Desalted peptides were finally dried in a speed vacuum and eluted in a solution C (2% FA in water).

### LC–MS/MS measurements

Peptides were centrifuged for 10 min at 18000g. Samples were measured using an Orbitrap Exploris 480 mass spectrometer (Thermo Fisher Scientific, San Jose, CA, United States) coupled to an UltiMate 3000 UHPLC system (Thermo Fisher Scientific, San Jose, CA, United States) equipped with a μ-precolumn (C18 PepMap100, 5 μm, 100 Å, 5 mm × 300 μm, Thermo Fisher Scientific, San Jose, CA, United States) and a homemade analytical column (Agilent Poroshell 120 EC-C18, 2.7 μm, 50 cm × 75 μm). 60 % of the total peptide solution was loaded in buffer A (0.1% FA in water) with a flow rate of 30 μL/min and eluted using a 90 min gradient at 300 nL/min flow rate. The gradient proceeds as follows: 9% solvent B (0.1% FA in 80% acetonitrile, 20% water) for 1 min, 9–13% for 1 min, 13–44% for 65 min, 44–55% for 5 min, 55–99% for 3 min, 99% for 5 min, and finally the system equilibrated with 9% B for 10 min. MS data were acquired in data-dependent acquisition (DDA) mode. The electrospray voltage was set at 2000 V, and the ion transfer tube temperature was set to 275°C. The full scan MS spectra were acquired at a resolution of 60,000 within the m/z range of 375–1600 using a “Standard” pre-set automated gain control (AGC) target. The RF lens was set to 40%, and the dynamic exclusion time was set to 14 s. For the MS2 acquisition, high-energy collision dissociation was performed with 28% normalized collision energy at an Orbitrap resolution of 15.000. Multiply charged precursor ions starting from m/z 120 were selected for further fragmentation. The AGC target was set to “Standard” and a 1.4 m/z isolation window was used for fragmentation.

### LC-MS/MS data analyses

LC-MS/MS Raw files were analyzed with MaxQuant (version 1.6.2.6) (Cox and Mann et al., 2008) for protein identification and label-free quantification (LFQ) using its integrated andromeda search algorithm against the reference proteome of Homo sapiens (UP000005640, number of entries: 82.485 (downloaded in April 2022). The match between runs feature was enabled and the search was performed allowing for fixed modifications; Carbamidomethyl (C) and variable modifications Oxidation (M); Acetyl (Protein N-term). The mqpar.xml file is provided alongside the deposited proteomics data. Using Perseus (version 1.6.14.0) (Tyanova et al., 2016), reverse hits, hits identified only by site and common contaminants were removed from the datasets at a false discovery rate (FDR) of equal or below 0.01. LFQ intensities were log_2_ transformed and only proteins found in at least three replicates (out of 4 samples) in the same group, were maintained for downstream analyses. Missing values were imputed from a normal distribution (1.8 standard deviations left-shifted Gaussian distribution with a width of 0.3 in relation to the standard deviation of measured values). Statistically enriched proteins were determined by a two-tailed *t*-test (FDR = 0.01, s0 = 0.1; l*og2* FC> 1.5; *-log10 p*-value >1.5, between the quadruplicates (MCPH1-BRCT-NLS-Turbo over NLS-Turbo control). Data were visualized using R (version 4.3.0). Proteins with a Fold Change (FC) equal or more than 1.5 and *-log10 p*-value >1.5 were considered as significantly enriched.

### Functional gene enrichment and network analysis

The lists of genes associated with specific functional categories were retrieved using AmiGO (http://amigo.geneontology.org/amigo), and from from The Gene Ontology (GO) Consortium (The Gene Ontology Consortium et al., 2023) on “09-01-2024”. The DDR repair list was assembled by combining genes retrieved from The Gene Ontology Consortium and from manually curated references (Lange et al., 2011). Gene Ontology term enrichment analysis was implemented with R using the “enrichplot” package using the total number of detected variables as a background. Heatmaps were implemented with the “ggplot2” package in R with a complete Euclidean hclust. Significantly enriched proteins identified in at least one of the six conditions (a total of 283 hits) were used for constructing a network using the Stringdb database (Szklarczyk et al, 2023). Visualization was based on hits associated with top Cellular Component GO terms. STRING interactions were added as links between proteins, with a cutoff set at a medium confidence (0.400). The Background for GO analyses comprised the total number of variables obtained amongst all the conditions. Hits associated to DDR (GO:0006974) were kept in the network. The network was retrieved from STRING. The presence of each protein in a defined condition was represented by a color-coded pie chart which was manually implemented.

## Acknowledgements

We thank members of the Baubec laboratory for their input and criticism. Furthermore, we thank the Biology Imaging Center at Utrecht University, the Utrecht University FACS facilities at the Science and Veterinary Faculties, and the Cell Microscopy Core at UMC Utrecht for their support. We thank Dr. Susanne MA Lens (University Medical Center Utrecht, Netherlands) for kindly providing the Human U2OS cells and prof. Dr. Wouter de Laat (Hubrecht Institute, Utrecht, Netherland) for sharing SA1 and SA2 antibodies. This work was supported by the European Research Council (865094 - ChromatinLEGO - ERC-2019-COG), Utrecht University, and the EMBO Young Investigator program. M.C.T.S and K.E.S. acknowledge support from the Horizon 2020 program INFRAIA project Epic-XS (Project 823839), and the NOW funded Netherlands Proteomics Center through the National Road Map for Large-scale Infrastructures program X-Omics (Project 184.034.019).

