## Supplemental Figures and Info for "Probing DNA damage sites reveals context-dependent and novel DNA damage response factors"

**Supplementary Information**

- **Supplementary Figures and Legends 1-10**
- **Supplementary Video Legends 1-6**
- **Supplementary Table Legends 1-2**

### Supplementary Figure Legends

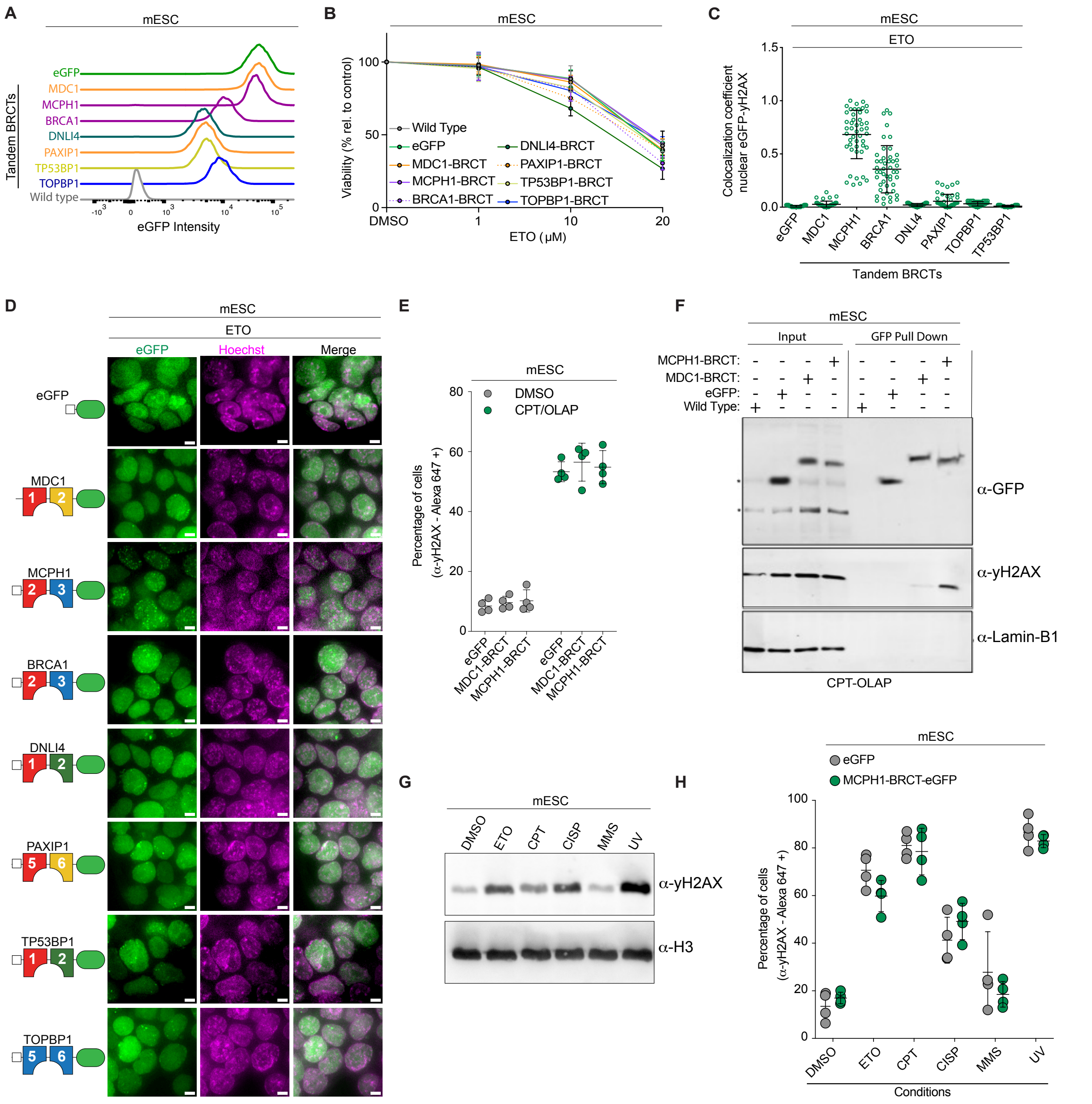


#### Figure S1 - Validation of eCRs against yH2AX in mESCs

**(A)** Representative FACS profiles showing stable and homogenous expression of selected tandem BRCTs fused to eGFP and eGFP alone in mESCs. **(B)** FACS Cell viability assays showing comparable viability among mESCs stably expressing the selected tandem BRCTs as compared to cells expressing nuclear eGFP and wild-type cells, following 12 h treatment with increasing concentrations of ETO. The viability percentage is relative to cells treated with DMSO control. Error bars represent ±SE of the mean of ten technical replicates. **(C)** Mander's correlation coefficient shows the colocalization coefficients of selected BRCT domain fused to eGFP relative to yH2AX sites. A minimum of 40 nuclei were analyzed and bars represent mean ±SD. Comparable results were obtained in three independent experiments. **(D)** Representative live-cell images of mESC stably expressing the indicated BRCTs fused to eGFP and the nuclear eGFP control. Cells were treated with ETO for 1h, stained with Hoechst, and immediately imaged. The scale bar is: 5 µM. **(E)** FACS-based quantification of yH2AX via anti-yH2AX antibody coupled to Alexa-Fluor 647 in the indicated mESCs treated with CPT+OLAP or with DMSO for 1h. Data represent the percentage ±SD of three biological replicates. Comparable results were obtained in one additional independent experiment. **(F)** Pulldown of the indicated eGFP fusion proteins in mESCs treated as in F. Wild-type cells were used as a control. Representative blots probed with antibodies against GFP, yH2AX, and Laminin-B1. Two independent experiments showed the same result. The asterisk denotes unspecific bands. **(G)** Western Blot analysis using histone acidic extracts obtained from wild type mESCs treated with the indicated agents for 1h. mESCs irradiated with UV (10 mJ/cm$^2$) were allowed to recover for 1h. Antibodies against H3 are used as a loading control. **(H)** FACS-based quantification of yH2AX via anti-yH2AX antibody coupled to Alexa-Fluor 647 in wild-type mESCs treated as in H. Data represents the percentage ±SD of three biological replicates. Comparable results were obtained in one additional independent experiment.

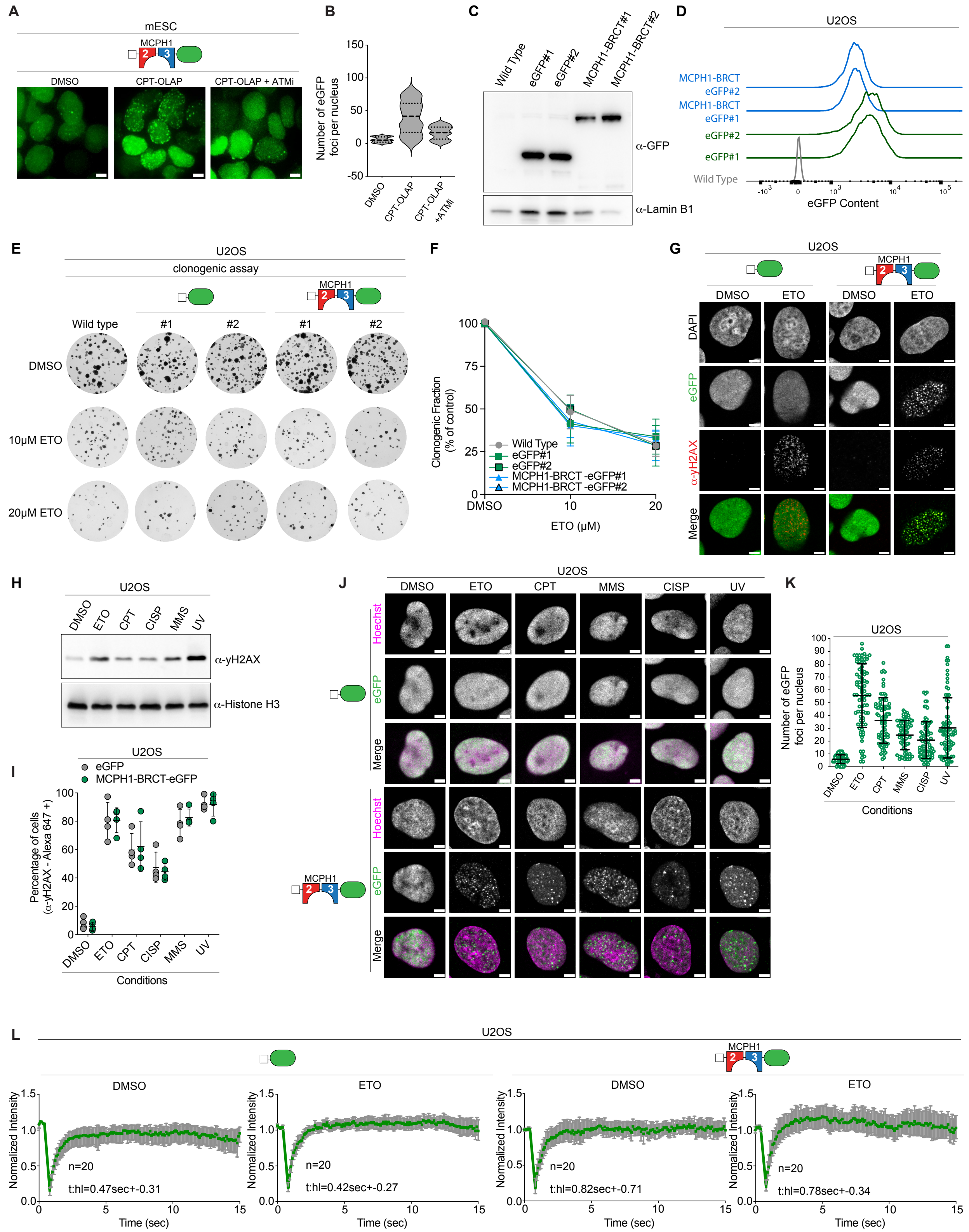

### Figure S2 - Functional Evaluation of MCPH1-BRCT-eCR

**(A)** Representative live-cell image of mESCs expressing MCPH1-BRCT-eGFP. Following treatment of mESCs with ATM inhibitor for 24h, cells were treated with CPT-Olap for 1h and immediately imaged. Scale bars: 5 μM. **(B)** Quantification of the number of discernible eGFP foci per nucleus from mESCs from A. Data represent mean (dashed thicker line) ±SD from at least 80 nuclei (segmented on the basis of eGFP background). Similar results were obtained in two additional experiments independently performed. **(C)** Western blot analysis comparing protein levels of two independently derived clones (#1 and #2) in nuclear extracts obtained from U2OS cells stably expressing the indicated eGFP constructs. Laminin B1 is used as a loading control. This experiment was performed two times with comparable results. **(D)** FACS profiles measuring the stable and homogenous expression of the independent U2OS clones described in C. **(E)** Clonogenic survival assays (crystal violet-stained plates) of independently derived U2OS clones treated as indicated for 24h. Subsequently, cells were incubated in fresh medium without the drug for an additional 15 days prior to staining. **(F)** Quantification of the signal intensity of plates from E, (normalized to the background) and relative to control DMSO cells, is shown. Data are presented as the mean SD; n=12. (**G**) Representative immunofluorescence images of U2OS cells (clones#1 for both cell lines) showing the nuclear localization of MCPH1-BRCT-eGFP and its colocalization with yH2AX (red). Cells expressing nuclear eGFP (clone#1) showed a diffuse nuclear signal. **(H)** Western Blot analysis using histone acidic extracts obtained from wild-type mESCs treated with the indicated agents for 1h. U2OS cells irradiated with UV (10 mJ/cm$^2$) were allowed to recover for 1h. Antibody against H3 is used as a loading control. **(I)** FACS-based quantification of yH2AX via anti-yH2AX antibody coupled to Alexa-Fluor 647 in wild type U2OS cells treated as in H. Data represent the percentage ±SD of three biological replicates. Comparable results were obtained in one additional independent experiment. **(J)** Representative images of fixed U2OS cells expressing MCPH1-BRCT-eGFP treated as indicated (see conditions shown in I) for 1h. Cells were fixed, stained with Hoechst (magenta), and immediately imaged. **(K)** Quantification of the number of microscopically discernible eGFP foci per nucleus from live U2OS cells from J. The data represent mean ±SD from at least 100 cells. Comparable results were obtained in two additional independent experiments. **(L)** FRAP recovery curves of U2OS cells stably expressing MCPH1-BRCT-eGFP or eGFP and treated as indicated for 1h. Data show the mean intensity (pre-bleached and background-normalized ROIS). Data represent mean ± SD from at least 20 ROIS in 20 randomly selected cells per condition. The t=hl (time of the half-life) of each curve as well as the number of ROIs are indicated. Two independent experiments showed comparable results.

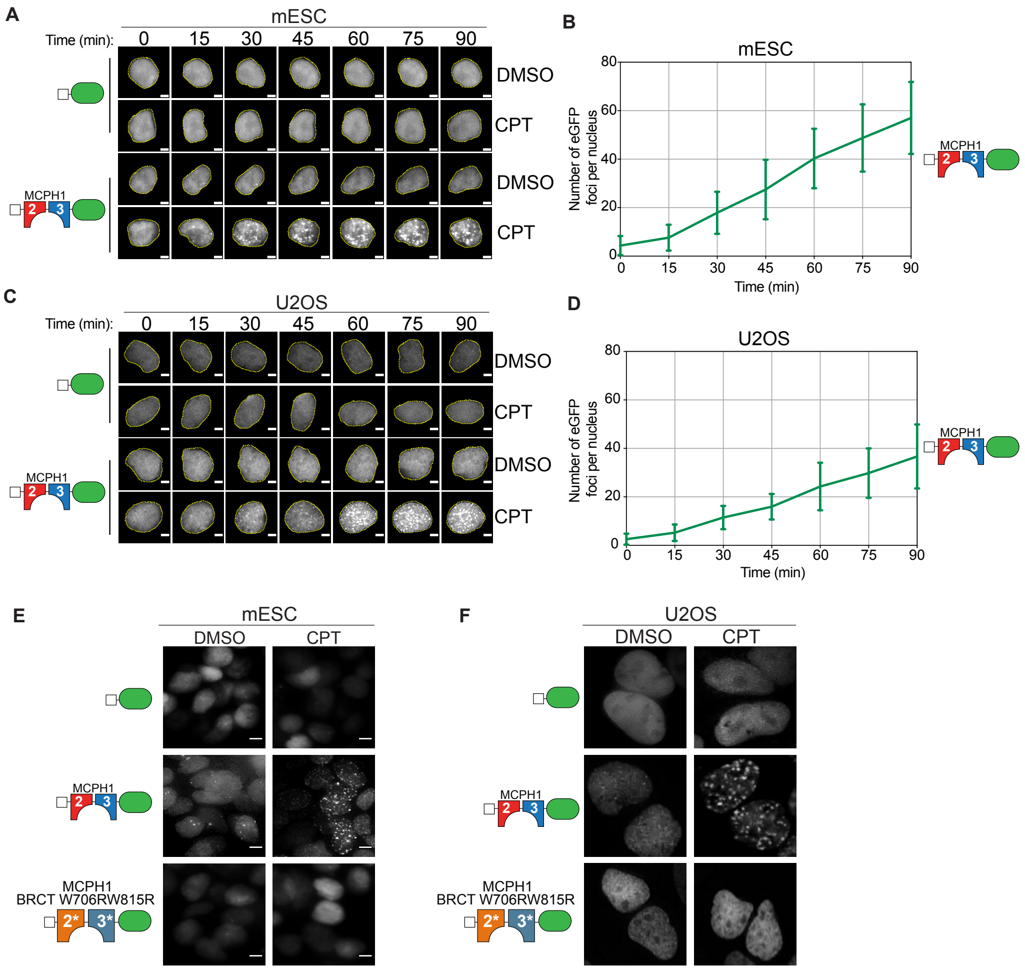


**Figure S3 - Using the MCPH1-BRCT eCR to track early yH2AX kinetics in living cells**

**(A)** Representative filmstrips of time-lapse fluorescence microscopy obtained from proliferating mESCs expressing MCPH1-BRCT-eGFP or nuclear eGFP from different time points and imaged after the indicated treatments. The ROI around selected nuclear eGFP is shown. **(B)** Quantification of nuclear eGP foci from mESCs described in A. Cells were imaged every 5 minutes right after the addition of the indicated agents and quantification was performed with at least 50 nuclei from two independent time-lapse microscopy experiments using the images captured every 15 min. **(C)** Representative filmstrips of time-lapse fluorescence microscopy obtained from proliferating U2OS cells expressing MCPH1-BRCT-eGFP or nuclear eGFP. **(D)** Quantification of microscopically discernible nuclear eGP foci from U2OS cells from C. Cells were imaged every 5 minutes right after the addition of the indicated agents and quantification was performed with at least 50 nuclei from two independent time-lapse microscopy experiments using the images captured at every 15 min. Representative live cell images of **(E)** mESCs or **(F)** U2OS cells showing the impaired recruitment of MCPH1-BRCT-eGFP due to the loss-of-binding point mutations (W706RW815R) in cells treated with CPT. Two independent experiments showed the same results. All scale bars are 5 μM.

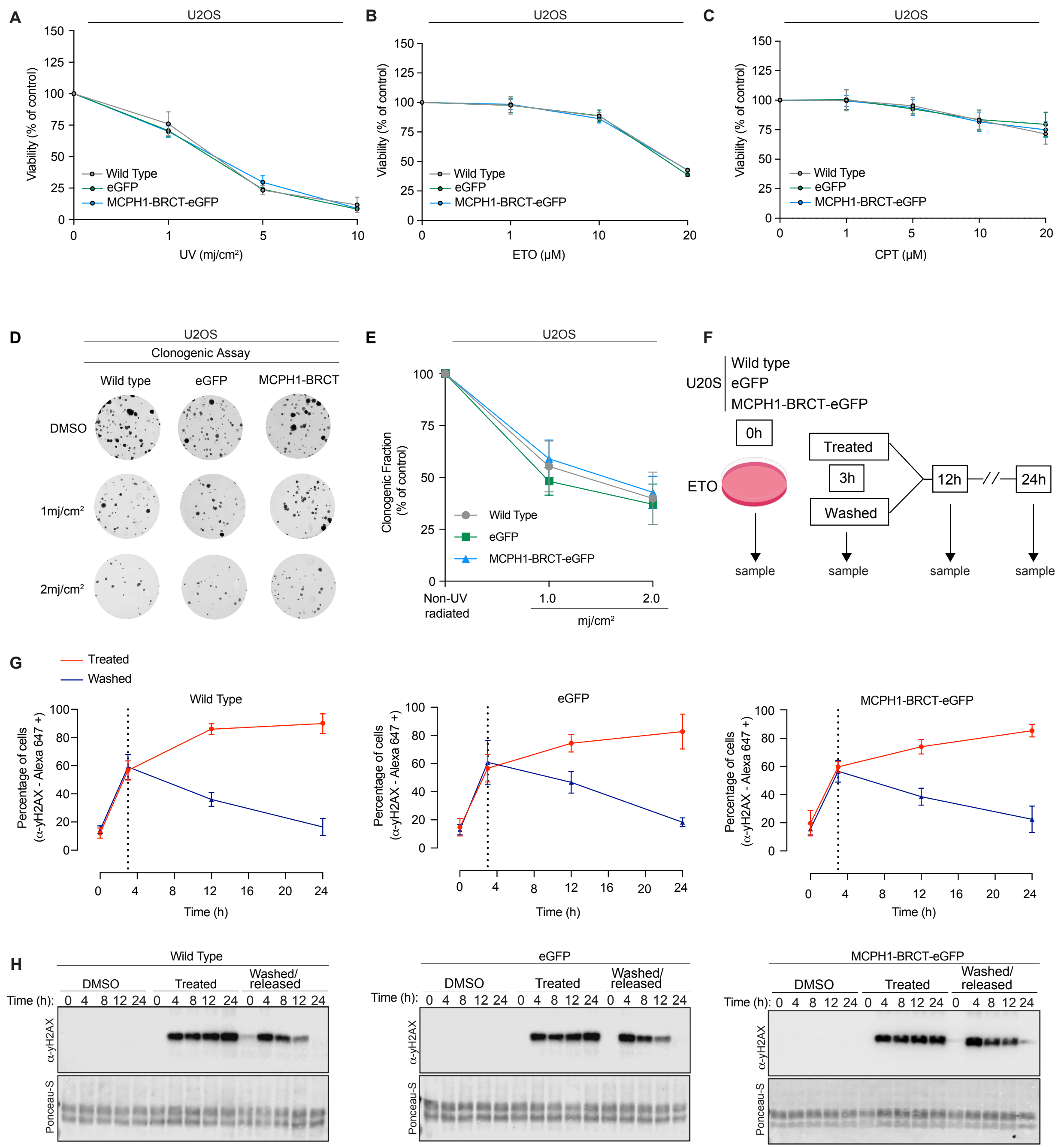


**Figure S4 - Expression of MCPH1-BRCT does not affect global DNA repair and cell viability**

**(A)** Plots showing comparable cell viability of U2OS cells recovering from the indicated UV irradiation conditions for 24h. Viability percentage was calculated relative to non-UV irradiated cells. Viability was measured after exposure of cells with **(B)** ETO or **(C)** CPT, as indicated, for 3h and recovery in fresh medium for an additional 12h. For plots in B and C, the percentage of viable cells was calculated relative to cells treated with DMSO control. Error bars represent ±SE of the mean of at least ten technical replicates. Experiments were independently performed two times. **(D)** Representative clonogenic survival assays of the indicated U2OS cells. Following UV irradiation, as indicated, cells were incubated in fresh medium for an additional 15 days prior to staining with crystal violet. **(E)** Quantification of the signal intensity per plate normalized to the background (region on the plate devoid of colonies) and relative to non-UV irradiated control cells showing comparable recovery from UV irradiation amongst the three different cell lines. Data are presented as the mean SD; n=9. Experiments were performed in duplicates. **(F)** Schematics - DNA damage recovery assays (Washout and release assay). Following exposure of the indicated U2OS cell lines with ETO for 3h, cells were either incubated in fresh medium without the drugs or kept in the same medium (Treated). Samples were collected at the indicated time points. **(G)** FACS-based quantification of yH2AX coupled to Alexa-Fluor 647 in samples collected at different time points showing comparable yH2AX levels in U2OS cells expressing MCPH1-BRCT-eGFP relative to eGFP control and wild-type cells in cells released in fresh medium (blue line). Data represent the percentage of positive cells ±SE of three biological replicates performed independently two times. Representative images of flow cytometry gates are shown in Figure S5A. **(H)** Representative Western Blots using acid histone extracts obtained from U2OS cells from experiments depicted in F and G and samples collected at intermediate intervals probed with antibodies against yH2AX. Histones detected on Ponceau-S were used as a loading control. The experiment was repeated twice.

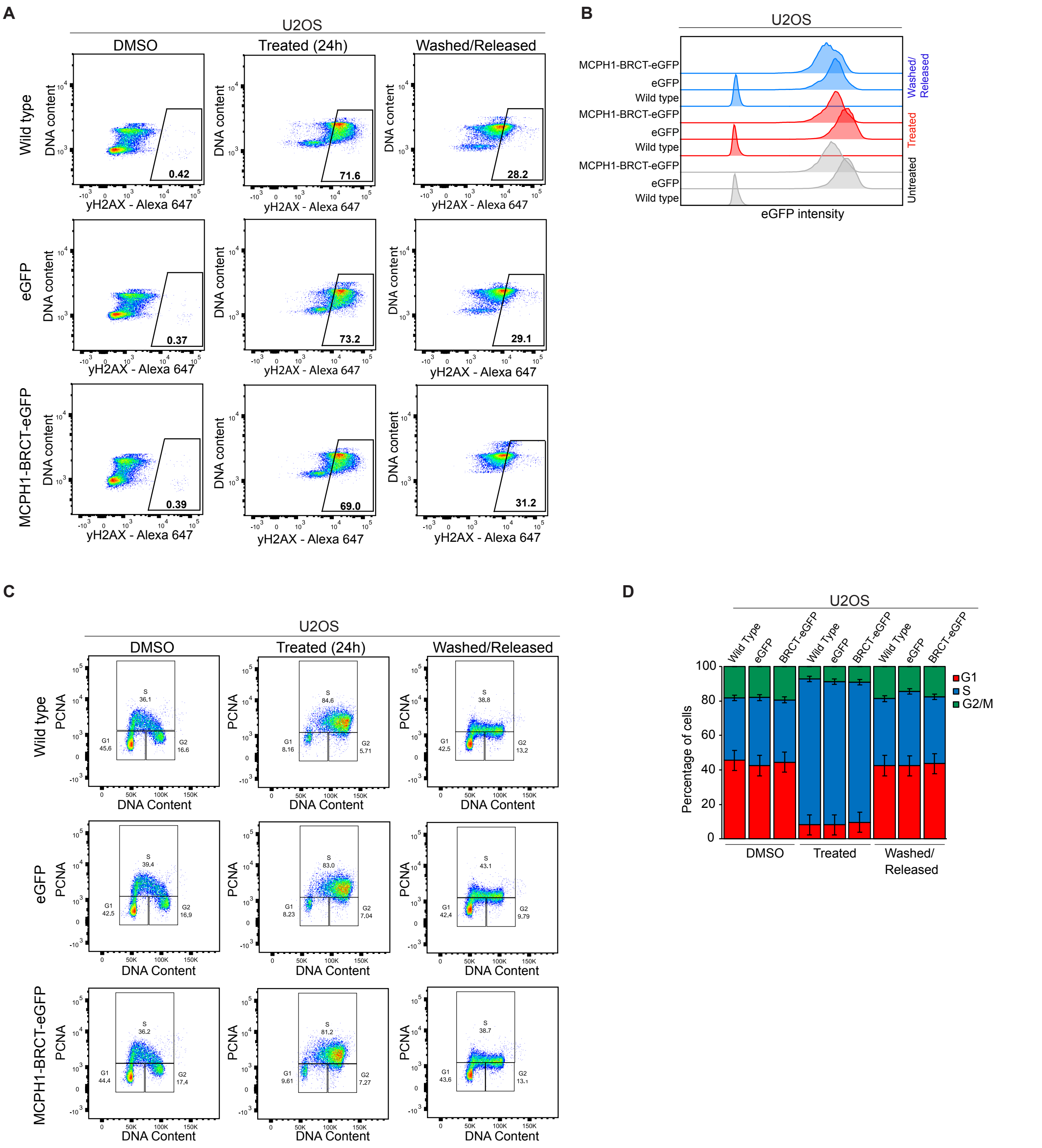


### Figure S5 - Expression of MCPH1-BRCT does not affect cell cycle progression

**(A)** Representative FACS profiles of yH2AX quantification in U2OS cells used in the Figure S4G. Values show the percentage of Alexa-Fluor 647 positive cells (*x-axis*). Cells were co-stained with DAPI (*y-axis*) to allow the simultaneous assessment of cell cycle progression. The experiment was repeated two times. **(B)** Representative FACS histograms showing the homogeneous expression levels of MCPH1-BRCT-eGFP and eGFP control in U2OS cells used in washout and release assays from Figure S4G. Data shows samples collected at 24h (see schematics in Figure S4F). **(C)** Representative scatter plots showing the DNA content distribution of cells measured by FACs. Cells were treated as depicted in Figure S4G (U2OS cells collected 24h after treatment and release in fresh medium). Cells were co-stained with PCNA to allow the measurement of cells in S phase and with DAPI. The relative distribution of cells in G1, S, and G2/M was based on specific controls of U2OS cells synchronized in G2/M and S phases with specific drugs. **(D)** Estimation of the relative distribution of U2OS cells in G1, S, and G2/M from C. Data represent the percentage ±SE of three biological replicates performed independently, two times.

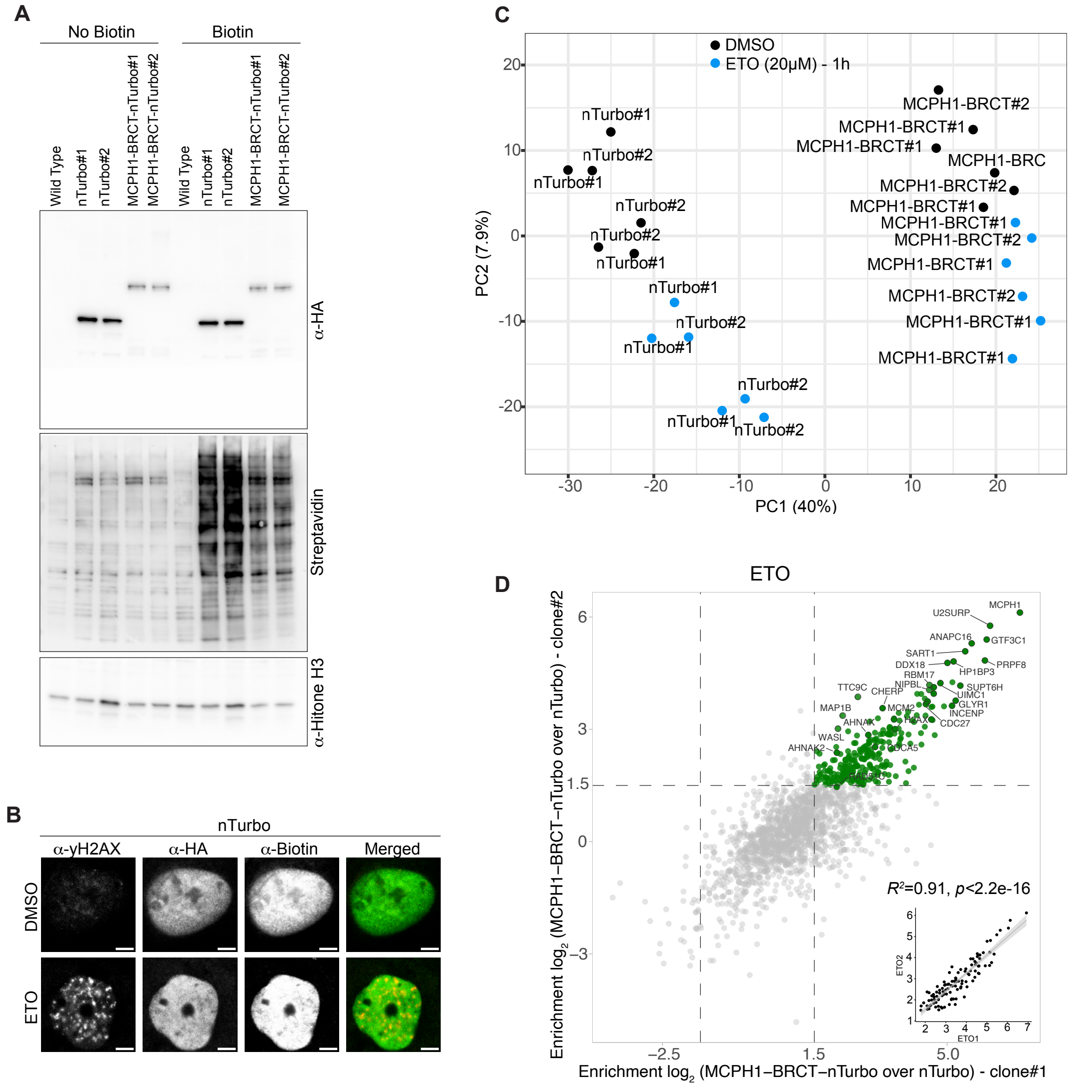


### Figure S6 - Functional validation of individually derived clones expressing TurboID constructs

**(A)** Western blot analyses of nuclear extracts obtained from two independently derived U2OS clones expressing either MCPH1-BRCT-Turbo or the nuclear Turbo (clones#1 and clones#2). Biotin treatments were done for 1h. Both clones present comparable levels of TurboID expression and global biotinylation as assessed by probing the membrane with Streptavidin-coupled to Alexa-Fluor 790 and antibodies against HA, respectively. Antibody against H3 was used as a loading control. Independent experiments were performed three times with comparable results. **(B)** Representative immunofluorescence images showing the localization of nuclear TurboID in U2OS (clone#1). nTurbo shows a diffuse nuclear distribution in cells co-incubated with biotin and with the indicated agents for 1h. Fixed cells were co-stained with antibodies against yH2AX, HA, and Biotin. This experiment was performed two times with similar outcomes. Scale bars: 5 µM. **(C)** Principal component analysis (PCA) plot of LFQ intensities from all the individual replicas (n=3 per condition) obtained from independently derived clones (MCPH1-BRCT-Turbo clones #1 and #2 and nTurbo clones #1 and #2) generated from TurboID experiments performed in U2OS cells co-incubated with either DMSO or ETO and biotin for 1h. **(D)** Scatter plot showing the $\log_2$ FC (LFQ intensities) of protein hits obtained through the direct comparison of the two independently derived clones (MCPH1-BRCT-Turbo#1 over nTurbo#1 versus MCPH1-BRCT-Turbo#2 over nTurbo#2) from U2OS cells co-incubated with ETO and biotin for 1h. Significantly enriched hits ($\log_2$FC>1.5) are shown in green. Manually selected hits are highlighted. The lower panel shows the Pearson correlation between the significantly enriched proteins from both clones ($R^2$:0.91), *P value <2.2.e-16*.

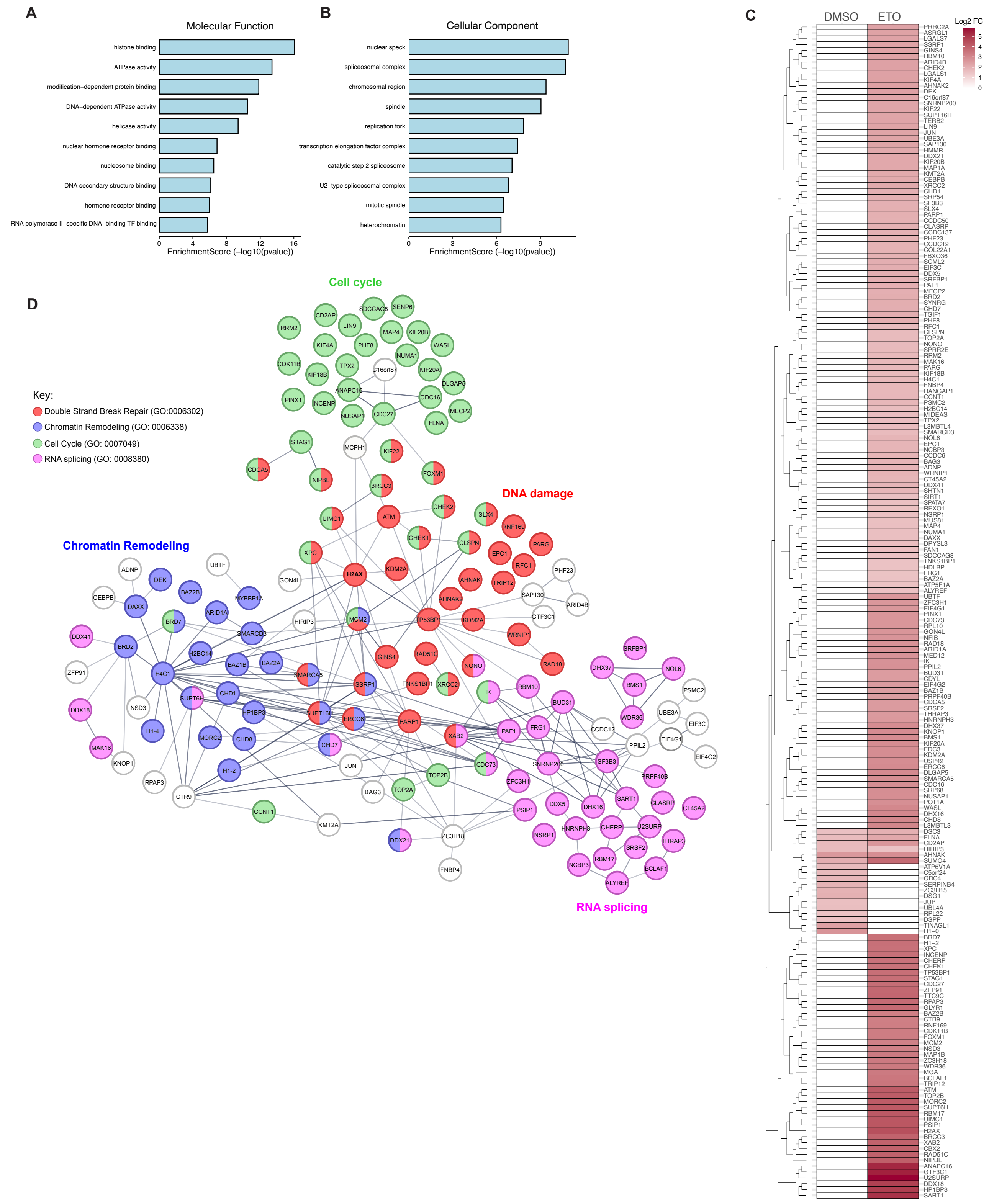


Legend on next page

**Figure S7 - Gene Ontology and and network maps of proteins associated to yH2AX sites triggered by ETO.**

**(A)** Bar plots showing the top ten Molecular Function GO terms associated with statistically enriched proteins (MCPH1-BRCT-Turbo/nTurbo after ETO treatments, $log_2FC>1.5$, n=189) based on enrichment score (-log10pvalue). **(B)** as in A, but for Cellular Component GO terms. **(C)** Heatmap visualization of statistically enriched proteins ($log_2FC>1.5$) identified by the MCPH1-BRCT-Turbo/nTurbo treated with DMSO (n=19) or ETO (n=189). The $Log_2FC$ of MCPH1 was not included. Proteins not determined in each condition are shown as white boxes. **(D)** Protein interaction network of statistically enriched proteins (MCPH1-BRCT-Turbo/nTurbo treated with ETO, $log_2FC>1.5$, n=189) grouped by statistically significant GO term annotations for biological process ranked by the number of observed gene counts in each GO-associated term. The total number of detected variables was used as background. Individual proteins are shown as nodes, colored on the basis of their associated GO-term (see key). Edges indicate interactions (score >0.4).

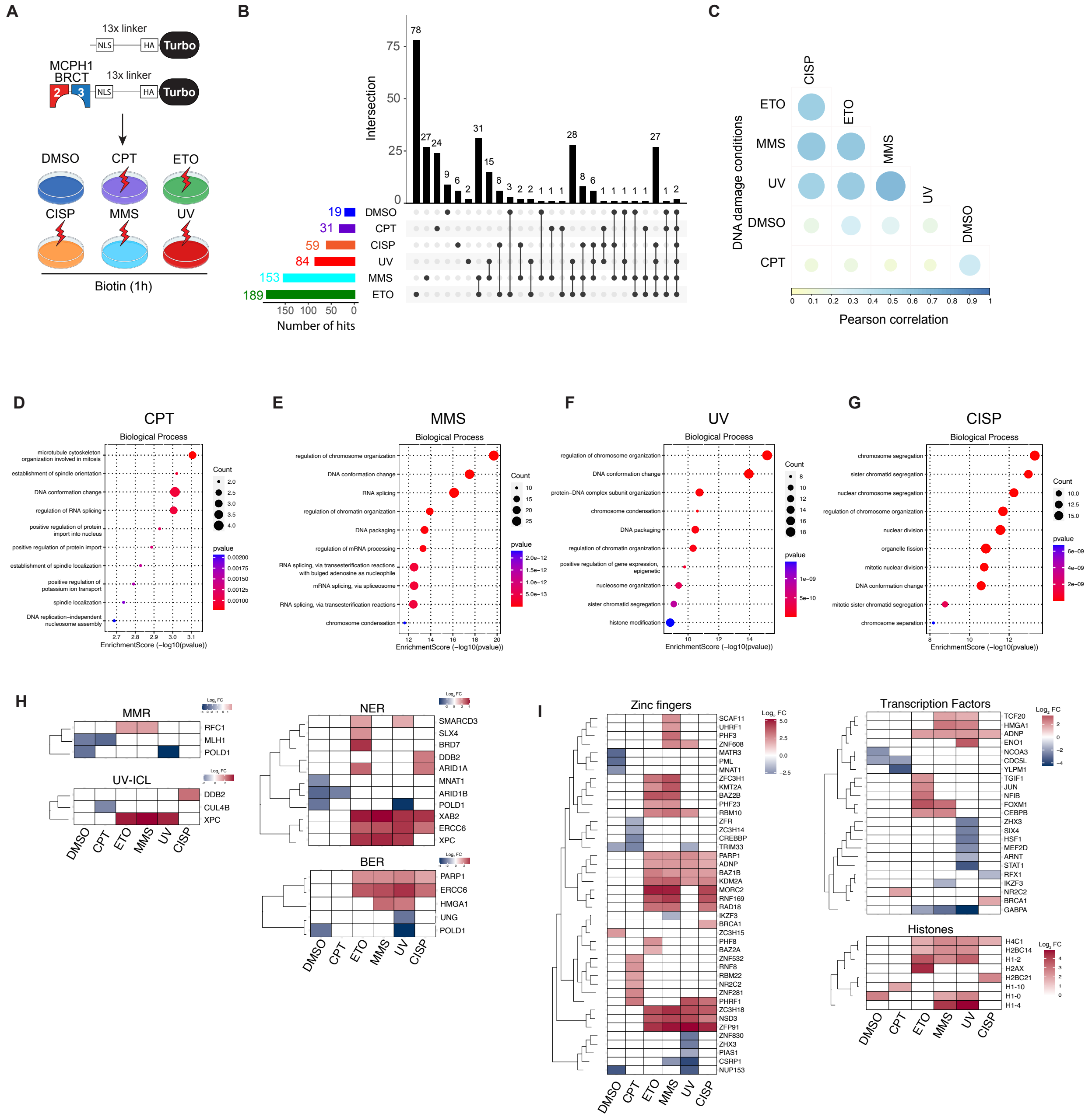


**Figure S8 - MCPH1-BRCT probes reveal the protein composition at DNA damage sites induced by different genotoxic agents**

**(A)** Schematics of DNA damage conditions used for TurboID experiments. (B) Upset plot showing the overlap of significantly enriched proteins (MCPH1-BRCT-Turbo/nTurbo) detected under the indicated conditions. The number of statistically significantly identified proteins is indicated. **(C)** Chart showing the Pearson correlation ($log_2FC$ based) among the different conditions. Circle sizes reflects the $R^2$ Pearson correlation. Cells treated with CPT show the lowest correlation relative to other conditions. **(D)** Bubble chart showing the top enriched Biological GO terms, ranked based on P-values, associated with statistically significant enriched proteins (MCPH1-BRCT-Turbo over nTurbo) treated with: **(D)** CPT, **(E)** MMS, **(F)** UV, and **(G)** CISP. The total number of variables detected in each condition was used as background. Heatmap visualization of statistically significantly identified proteins ($Log_2FC>1.5$) classified according to different functional categories in DDR **(H)** and other functional categories **(I)**. Proteins were clustered according to their LFQ ($Log_2$ FC), obtained under the identified conditions. Proteins not determined in each condition are shown as white boxes.

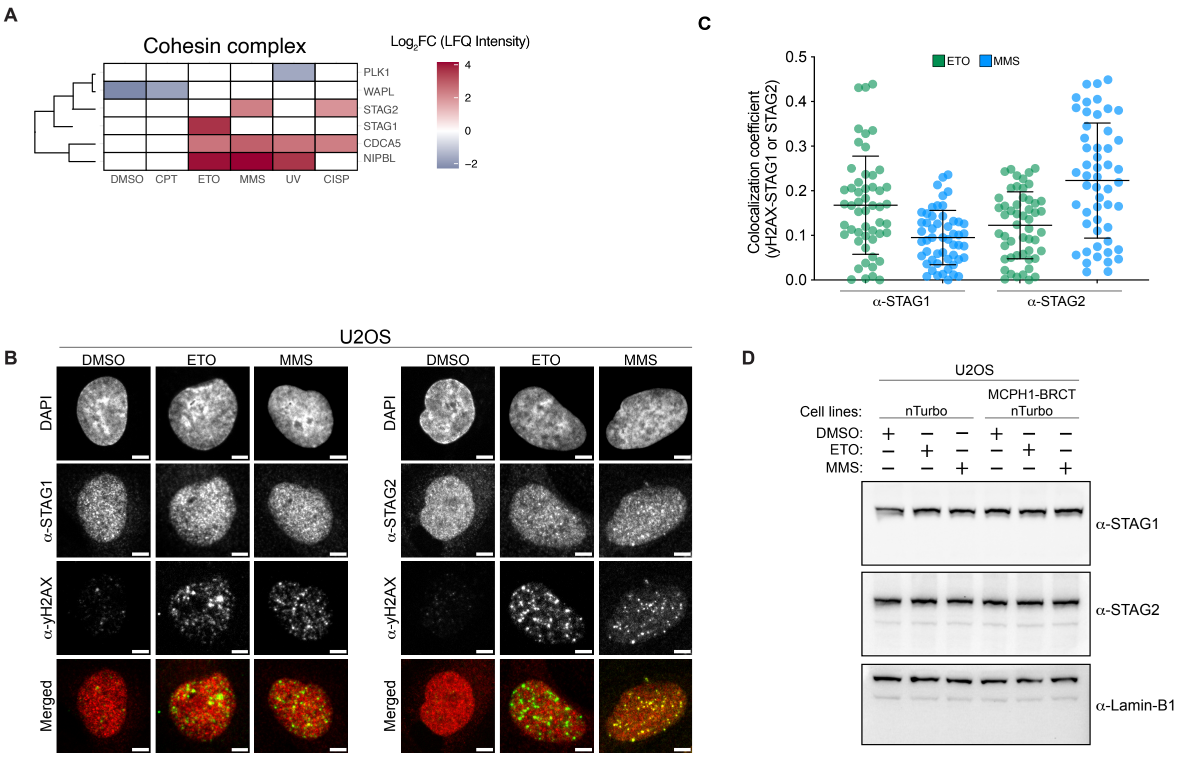


**Figure S9 - Recruitment analyses of STAG1/STAG2 to damaged chromatin reveal increased association of STAG2 to MMS- induced yH2AX sites**

**(A)** Heatmap visualization of members of the cohesin complex statistically significantly identified ($Log_2FC>1.5$) under the indicated treatments. **(B)** Representative immunofluorescence images of fixed wild type U2OS cells showing the nuclear localization of STAG1, STAG2 (independently stained, red), and yH2AX (green) in cells exposed to the identified treatments for 1h. The scale bar is: 5 µM. **(C)** Quantification of colocalization of STAG1 or STAG2 with yH2AX was determined using Mander's colocalization coefficient. A minimum of 50 nuclei were analyzed and bars represent mean ±SD. Comparable results were obtained in two independent experiments. **(D)** Western Blot analysis showing comparable amounts of STAG1 and STAG2 in nuclear extracts obtained from the U2OS cell lines used in the proximity biotinylation assays, treated as indicated. Blots were probed with anti-STAG1, anti-STAG2, and anti-Laminin-B1 (used as a loading control). Comparable results were obtained in two independent experiments.

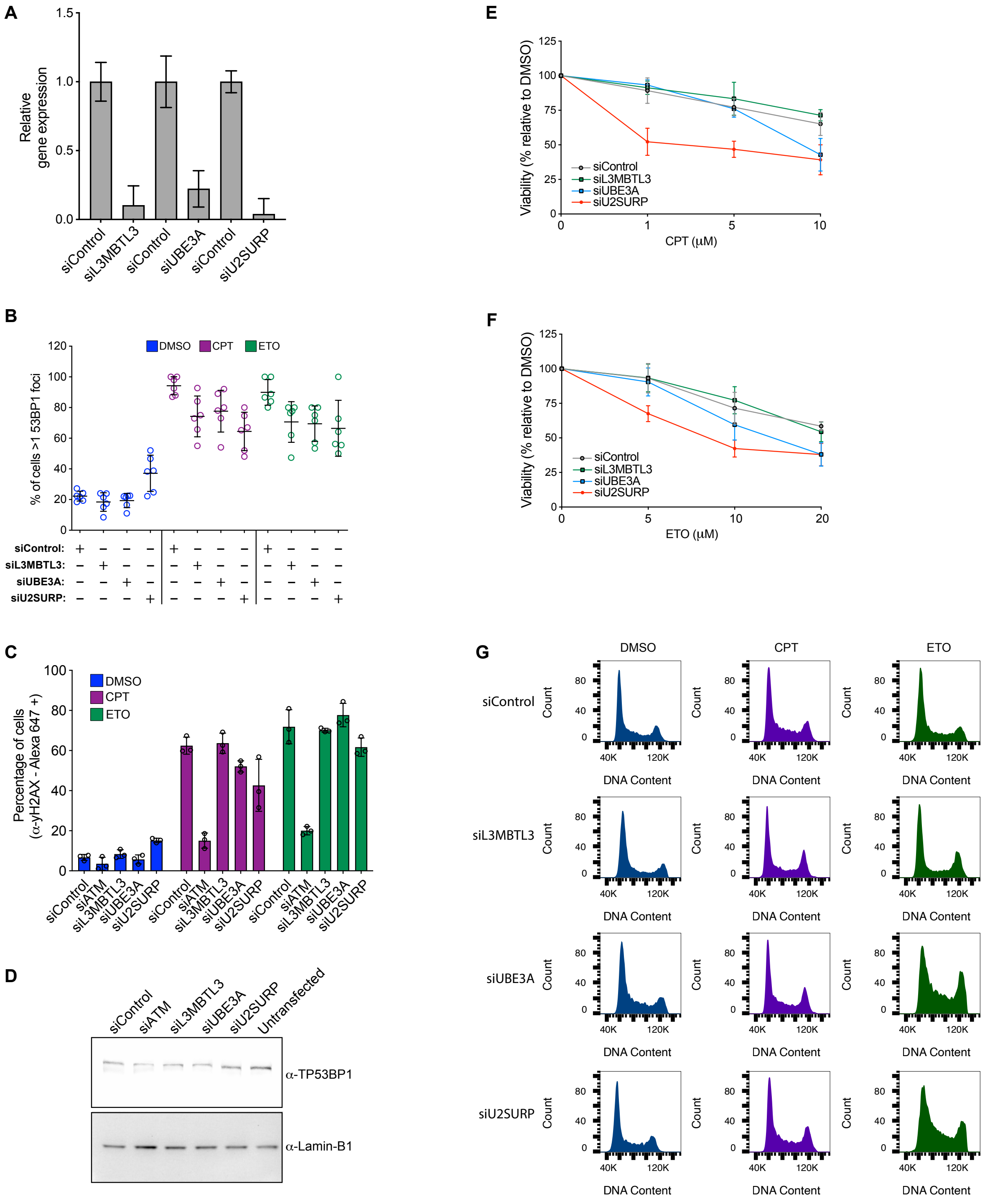

**Figure S10 - Effect of depletion of U2SURP, UBE3A and L3MBLT3 on DDR**

**(A)** Bar plots showing the depletion of the indicated mRNAs following siRNA transfection for 48h in unperturbed U2OS wild-type cells. Expression of each gene was measured by qPCR and normalized to GAPDH. Data represent the fold change (mean ± SE of three independent experiments performed in triplicate) relative to siRNA control (siControl). **(B)** Quantification of cells with more than three 53BP1 discernible foci after siRNA-mediated depletion for 48h in wild-type cells treated as indicated (images depicted in Figure 6A). Datapoints represent the mean ±SD from at least 40 cells in six different biological replicas. **(C)** FACS-based quantification of yH2AX via yH2AX antibody coupled to Alexa-Fluor 647 following siRNA-mediated depletion of the indicated proteins and treatment with the indicated drugs for 1h in wild type U2OS cells. Data represent the percentage of positive cells ±SE of three biological replicates. Comparable results were obtained in one additional independent experiment. **(D)** Western blot analysis comparing the levels of the protein 53BP1 in nuclear extracts obtained from wild type U2OS cells depleted for the indicated proteins for 48h. Untransfected cells were included as a control. Laminin-B1 is used as a loading control. This experiment was performed two times with comparable results. Cell viability assays of U2OS cells depleted as in A and treated with different drug concentrations of: **(E)** CPT or **(F)** ETO, as indicated, for 1h. Cells were then washed and incubated in fresh medium for 12h. The percentage of viability is relative to control cells (DMSO) and was measured using the CellTiter-Glo kit. Data represent mean ±SE of at least ten technical replicates from two independent experiments. **(G)** Representative FACS histograms of wild-type U2OS cells treated as indicated for 1h after siRNA-mediated depletion with the indicated siRNAs for 48h. The relative distribution of cells in G1, S, and G2/M is shown in Figure 6D.

**Supplemental Files Legends**

**Video S1**

Representative time-lapse fluorescence video of proliferating mESCs expressing MCPH1-BRCT. Cells were imaged at 5-min intervals for approximately 15 h post addition of medium containing DMSO. Images were processed according to methods. Scale bar, 50 μm.

**Video S2**

Representative time-lapse fluorescence video of proliferating mESCs expressing MCPH1-BRCT under DNA damage conditions. Cells were imaged at 5-min intervals for approximately 15 h post addition of medium containing CPT. Scale bar, 50 μm.

**Video S3**

Representative time-lapse fluorescence video of proliferating mESCs expressing nuclear eGFP control. Cells were imaged at 5-min intervals for approximately 15 h post addition of medium containing DMSO. Scale bar, 50 μm.

**Video S4**

Representative time-lapse fluorescence video of proliferating mESCs expressing nuclear eGFP control under DNA damage conditions. Cells were imaged at 5-min intervals for approximately 15 h post addition of medium containing CPT. Scale bar, 50 μm.

**Video S5**

Representative time-lapse fluorescence video of proliferating U2OS cells expressing MCPH1-BRCT. Cells were imaged at 5-min intervals for approximately 15h post addition of medium containing DMSO. Scale bar, 10 μm.

**Video S6**

Representative time-lapse fluorescence video of proliferating U2OS cells expressing MCPH1-BRCT under DNA damage conditions. Cells were imaged at 5-min intervals for approximately 15h post addition of medium containing CPT. Scale bar, 10 μm.

**Table S1**

List of mouse BRCT-containing proteins and the amino acid coordinates of the tandem BRCTs used as probes to target yH2AX, related to Figure 1.

**Table S2**

List of proteins identified by the DNA damage sensor MCPH1-BRCT in close proximity to yH2AX in U2OS cells exposed to DMSO, CPT, ETO, CISP, MMS and UV radiation.
